# Mechanical model of muscle contraction. 5. Tension rise after phase 1 of a length step

**DOI:** 10.1101/2019.12.16.878868

**Authors:** S. Louvet

## Abstract

The theoretical approaches developed in accompanying Papers 1 to 3 lead to calculations of the isometric tetanus tension (T0) and the minimum tension (T1) observed at the end of phase 1 of a length step where the fiber is shortened (see Paper 4). During the next three phases, the time rise of the tension (T), from T1 to T0, is determined for any step (see Supplement S5.K). The tension T is expressed as a master equation which is the sum of five terms: (a) T1, (b) a positive or zero contribution resulting from the relaxation induced by the disappearance of the viscosity forces present during phase 1, (c) a positive contribution of elastic origin resulting from the new myosin II heads initiating a working stroke (WS) in the blank areas, (d) a negative contribution caused by the fast detachment of the heads still strongly attached and whose orientation of the levers is beyond the *up* position, (e) a negative contribution caused by the slow detachment of WS heads whose orientation of the levers is close to the *up* position. An agreement between the model equation and the experimental results referenced in the physiological literature is proven (r^2^>97.5%). The kinetics of each of the theoretical curves make it possible to distinguish phases 2, 3 and 4 characteristics of the tension rise to T0. The criteria defined to describe the tension at the end of phase 2 (T2) are applied to the master equation. There is an adequate adjustment between the theoretical and experimental T2 values for shortenings less than 8 nm in modulus (r^2^ > 97%).

## Introduction

An isolated muscle fiber is isometrically tetanized and the measured tension reaches a plateau (T0). The stimulated fiber is suddenly shortened by a length step (ΔL). Four phases are distinguished; see Table 1 in [1] or Table 4 in [2]. Phase 1, which is studied in accompanying Paper 4, represents the sudden drop in tension concomitant with the shortening of the fiber. The following three phases describe the rise in tension. There is a rapid rise for 1 to 2 milliseconds (phase 2), followed by a slowdown of a few milliseconds with the possible presence of a plateau (phase 3), and finally a slow rise up to T0 extending over several tens of milliseconds (phase 4).

**Table 1.** Reference values for parameters relating to muscle fibers isolated from vertebrate (frog) muscles.

|  | From Fig 11<br>in Ford 1977 | From Fig 12<br>in Ford 1977 | From Fig 2<br>in Piazzesi 1992 |
| --- | --- | --- | --- |
| Fig No. in<br>Paper 5 | 1 | 2 | 3 |
| Animal | Rana T <sup>(1)</sup> | Rana T <sup>(1)</sup> | Rana E <sup>(2)</sup> |
| Muscle | TAM <sup>(3)</sup> | TAM <sup>(3)</sup> | TAM <sup>(3)</sup> |
| $\Gamma$ | 2.5 °C | 1 °C | 4 °C |
| $\tau_{p1}$ | 200 μs | 200 μs | 120 μs |
| T0 | 245 kPa | 243 kPa | 195 kPa |
| $L_{fm}$ | 6 mm | 6 mm | 5.2 mm |
| $N_{hs}$ | 5500 | 5500 | 5000 |
| $L0_s$ | 2.2 μm | 2.2 μm | 2.1 μm |
| $\delta X_{pre}$ | 6 nm | 6 nm | 5.5 nm |
| $\delta X_{Max}$ | 11.5 nm | 11.5 nm | 11 nm |
| $\delta X_T$ | 8 nm | 8 nm | 8 nm |
| $\delta X_{z1}$ | 3.5/4.5 nm | 4//5 nm | 3/4 nm |
| $K_{z1}$ | 1.4 | 1.35 | 1.6 |
| $P_{startF}$ | 0.6/0.63 | 0.6/0.63 | 0.57/0.61 |
| $P_{startS}$ | 0.3 | 0.3/0.31 | 0.3 |
| $\tau_{Relax}$ | 0.3/0.5 ms | 0.3/0.5 ms | 0.3/0.5 ms |
| $\tau_{startF}$ | 0.6/0.8 ms | 0.6/0.8 ms | 0.6/0.8 ms |
| $\tau_{preS}$ | 2 ms | 2 ms | 2 ms |
| $\tau_{startS}$ | 27 ms | 27 ms | 27 ms |
| $\tau_{preVS}$ | 50 ms | 50 ms | 50 ms |
| $\tau_{startVS}$ | 70 ms | 70 ms | 70 ms |
| $\tau_{preFDE}$ | 1 ms | 1 ms | 1 ms |
| $\tau_{FDE}$ | 5 ms | 5 ms | 3 ms |
| $\tau_{preSDE}$ | 7 ms | 7 ms | 5 ms |
| $\tau_{SDE}$ | 20 ms | 20 ms | 15 ms |
| $r^2$ | > 97.5% | > 97.5% | > 98% |

<sup>(1)</sup> *Rana Temporaria*
<sup>(2)</sup> *Rana Esculenta*
<sup>(3)</sup> *Tibialis Anterior* Muscle

Several articles are devoted to the modelling of the tension rise from T1 to T0 during phases 2, 3 and 4. The answer is always given takes the form of an algebraic sum of exponentials whose coefficients are calculated, either empirically or using the kinetic coefficients relating to the reversible reactions of the cross-bridge cycle; see equation (5) and Fig 23 from [2], Fig 3 in [3], equations 1 to 3 and Figs 3A and 3D from [4], Fig 2 and Appendix from [5]; Fig 1 in [6]; Fig 14 in [7].

**Fig 1.**
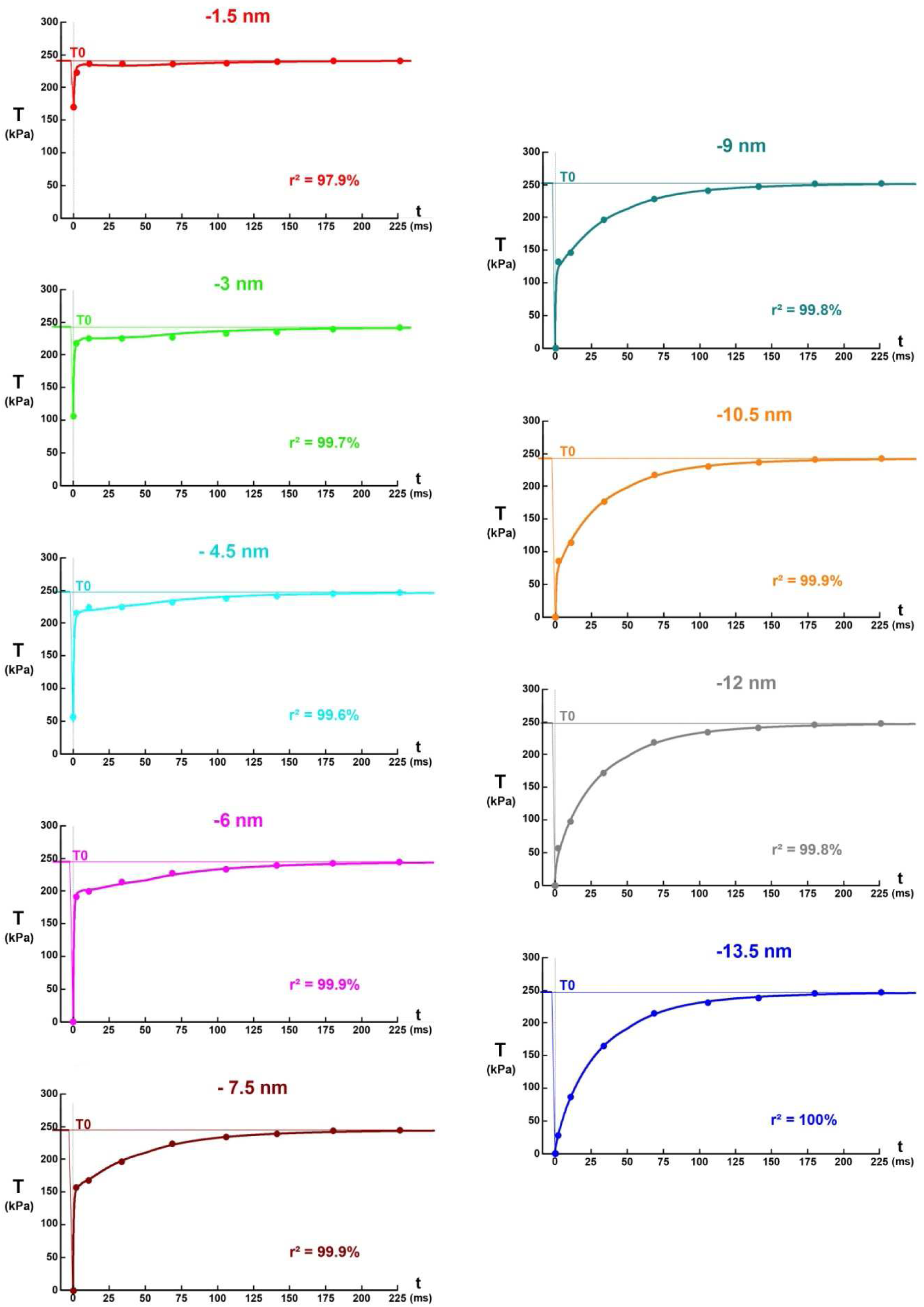
Instantaneous rise of the tension (T) up to the isometric tetanus plateau (T0). The theoretical curves are calculated from equation (1) for 9 length steps per hs varying from −1.5 nm to −13.5 nm. The points displayed on each of the 9 graphs are collected on the corresponding plots in Fig 11 from [2].

**Fig 2.**
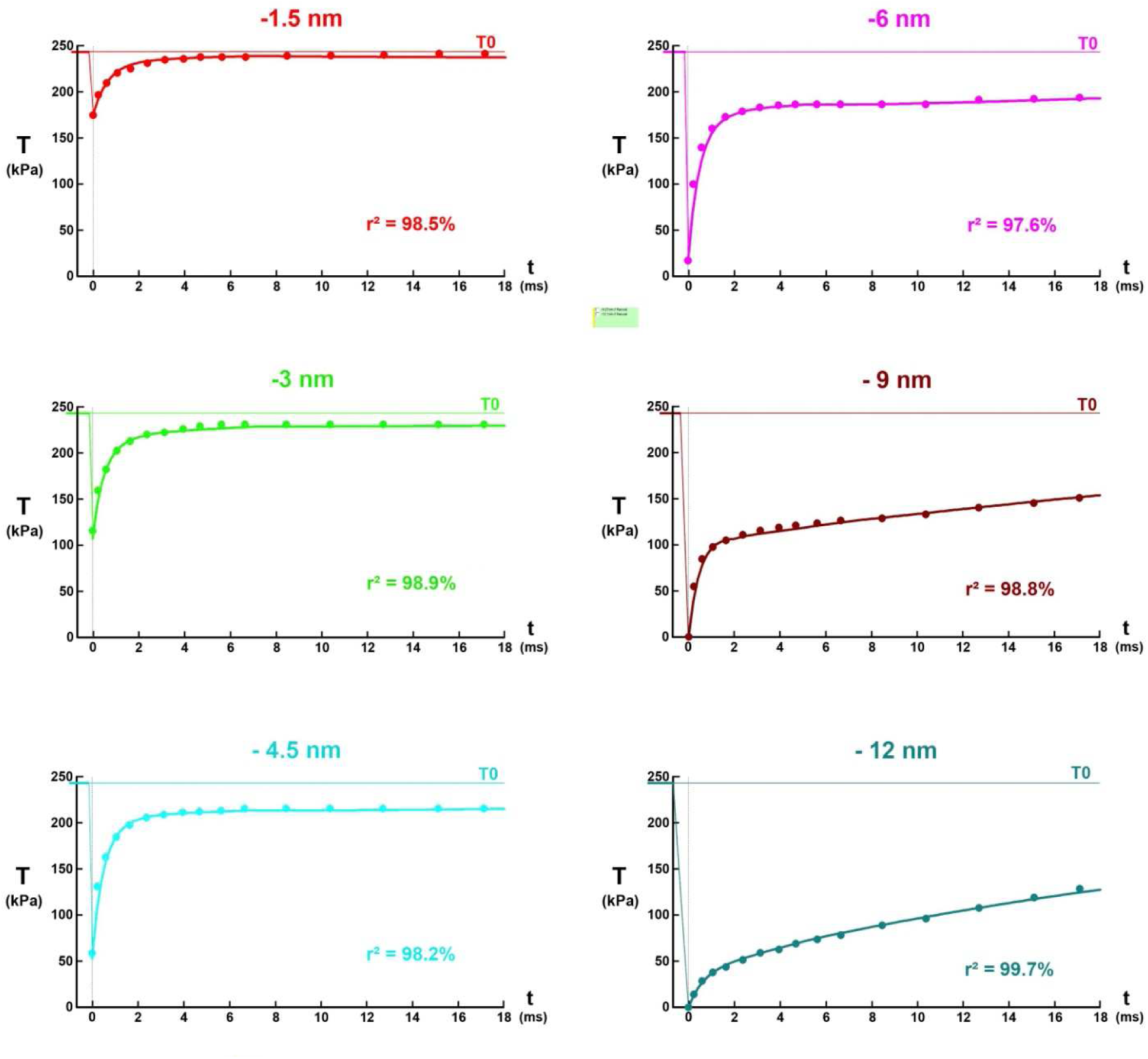
Instantaneous rise of the tension (T) during the first 18 ms. The master equation formulated in (1) is represented by a continuous line of different color for each of the 6 hs shortenings (ΔX = −1.5, −3, −4.5, −6, −9 and −12 nm). The colour points are taken from the 6 plots corresponding to each of the 6 length steps per hs in Fig 12 from [2].

**Fig 3.**
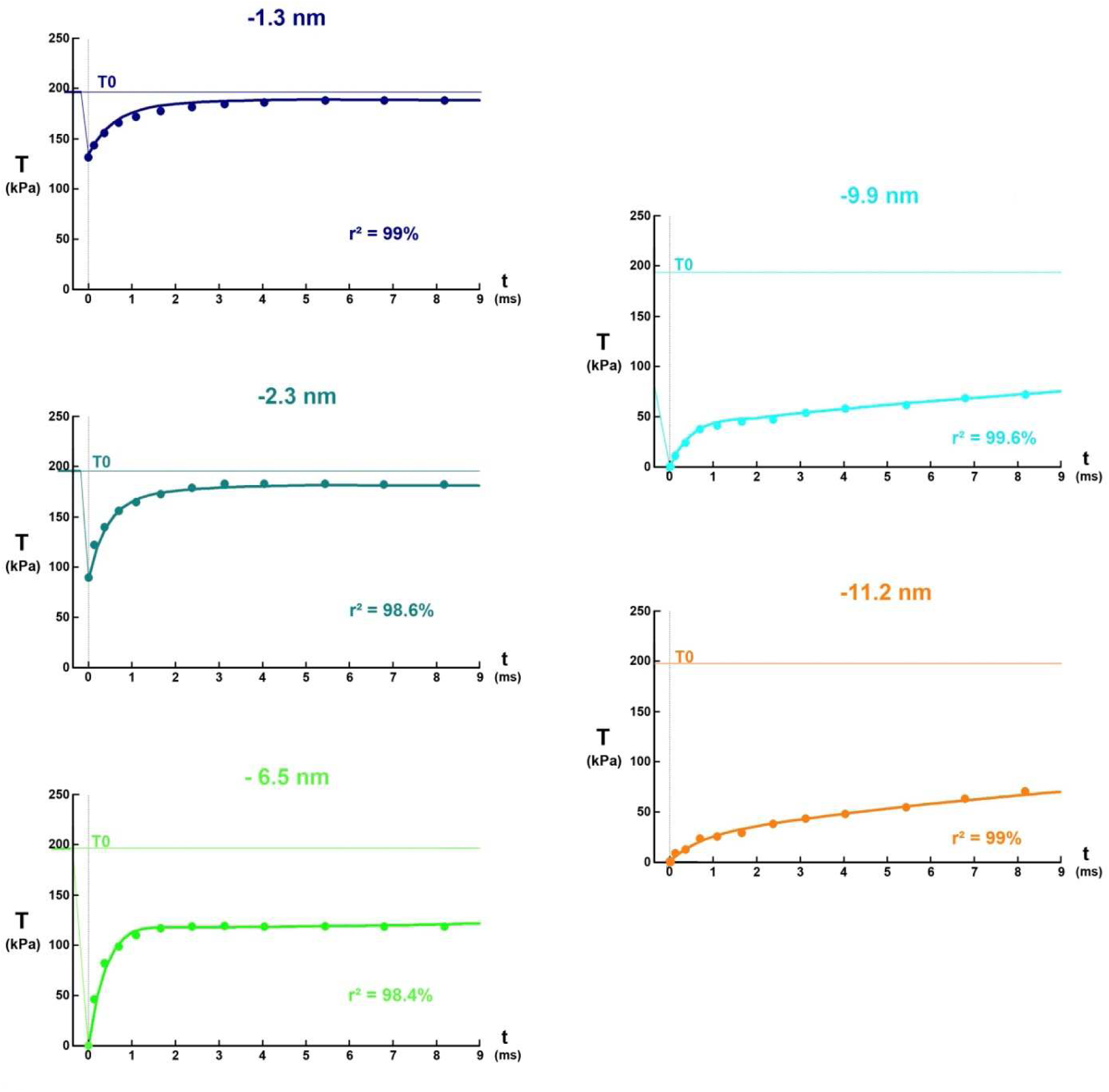
Fig 2. Instantaneous rise of the tension (T) during the first 9 ms. The master equation formulated in (1) is represented by a continuous line of color for each of the 5 hs shortenings (ΔX = −1.3, −2.3, −4.5, −6.5, −9.9 and −11.2 nm). The color points are collected from the 5 plots corresponding to the 5 length steps per hs in Fig 2 (left column) from [8].

We follow the same path by proposing a general equation that is also an algebraic sum of exponentials. However, this equation is based on the temporal parameters associated with cross-bridge cycle reactions assumed to be irreversible (see Hypothesis 1 in Supplement S1.B of Paper 1), and on the geometric and mechanical calculations for a myosin head and a half-sarcomere (hs) developed in accompanying Papers 2, 3 and 4.

## Methods

### Events before and after the Working Stroke

Hypothesis 4 presented in accompanying Paper 2 states that the lever of a WS myosin II head moves in a fixed plane, the lever orientation being defined by the angle θ in this plane. The maximum variation of θ (δθ_Max_) is equal to 70°, a range delimited by the two angles θ_up_ and θ_down_ relative to the two positions *up* and *down*. During the isometric tetanus plateau, the spatial density of θ is uniformly distributed over an angular range (δθ_T_) equal to about 50° and framed by the angles θ_up_ and θ_T_ (see Paper 4).

The 5 events briefly described below occur traditionally during the cross-bridge cycle (see supplement S1.B of Paper 1).

1/ The {WSstart} event generating the WS state is developed in paragraph B.3. According to hypothesis 2, it is divided into 3 modes, {startF}, {startS} and {startVS}, which refer, respectively, to a Fast, Slow and Very Slow initiation. When a head is in WS, it exerts a linking action on the actin filament and the myosin filament to which it is bound, an action that positively contributes to the value of the tension exerted across the hs.

2/ The state preceding the WS is the Strong Binding state described in paragraph B.4. In our model, the {startF} event particularizes the fast transition between Strong Binding and WS states.

3/ According to hypothesis 3, the event contributing to the detachment of a myosin head is divided into two modes, fast and slow.

The {FastDE} event is about the rapid detachment of WS heads whose lever has an angular position θ beyond θ_down_ after a length step. It is discussed in paragraph B.6.

The {SlowDE} event concerns the slow detachment of WS heads whose lever has an angle θ between θ_down_ and θ_T_ after a length step. It is explained in paragraph B.7.

When a head detaches according to {FastDE} or {SlowDE}, i.e. transits to the Detachment state, the tension it previously exerted disappears and this temporal abolition contributes negatively to the value of the tension of the hs and myofibril.

The irreversibility of the reactions preceding and closing the WS state allows the temporal equation of these 5 events, {startF}, {startS}, {startVS}, {FastDE} and {SlowDE}; see Supplements S1.A and S1.B of Paper 1. These various expressions are recalled in paragraph K.1 of Supplement S5.K.

### Master equation of the model

At the end of phase 1 of a length step, the fiber has shortened from ΔL and remains in isometry. After the effects of viscosity have disappeared, each half-sarcomere (hs) of the fiber is shortened by an identical length (ΔX) such that:

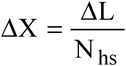

where ΔL is the value of the length step relating to the fiber, positive value for lengthening and negative value for shortening; N_hs_ is the number of hs per myofibril.

All the calculations inherent in establishing the master equation that governs the tension rise are provided in Supplement S5.K. The equation is formulated from:

- Proportions of heads initiating a WS fast or slowly (p_startF_, p_startS_)

- Time parameters associated with irreversible cross-bridge cycle reactions (τ_startF_, τ_preS_, τ_startS_, τ_preVS_, τstartVS, τpreFDE, τFDE, τpreSDE, τSDE)

- Geometric criteria specific to a myosin head and hs (X_down_, δX_E_, δX_pre_, δX_T_, δX_Max_)

The drafting of all the detailed terms that make up this equation requires at least two pages. To overcome this problem, a concise wording is proposed in the corpus of this section.

At the end of phase 1 (t=0), the relative tension (pT) is the sum of 5 terms:

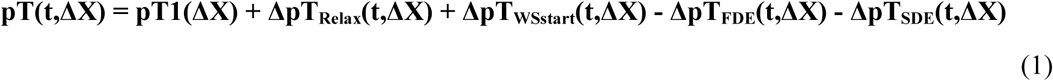

where **pT1** is the minimum tension at the end of phase 1 corresponding to time t=0. The determination of pT1 is the objective of Paper 4; its expression is recalled in equation (K29) to sub-paragraph K.5.1

**ΔpT**_**Relax**_ is the positive contribution due to the phenomenon of rapid relaxation caused by the disappearance of viscosity forces, the calculation of which is detailed in sub-paragraph K.5.2 and formulated in (K32)

**ΔpT**_**WSstart**_ is the positive contribution due to the new heads that initiate a WS in the areas freed by the shortening; the calculations are developed in paragraph K.2 in 4 domains and are grouped in equation (K34) delivered in sub-paragraph K.5.3

**ΔpT**_**FDE**_ is the negative contribution due to the rapid detachment of heads from the *down* state. It is evaluated in paragraph K.3 and expressed with (K35) in sub-paragraph K.5.4

**ΔpT**_**SDE**_ is the negative contribution due to the slow detachment of the WS heads whose levers have an θ angle between θ_up_ and θ_T_. It is explained in paragraph K.4 in 6 sectors. Its formula is given in (K37) to sub-paragraph K.5.5

### Objective

The purpose of the paper is to compare equation (1) with the experimental curves presented in the physiological literature.

### Algorithmic

The sequence of computer programs is written in Visual Basic 6. Equation (1) was developed in algorithmic form to obtain plots of relative tension (pT) as a function of time t for different values of ΔX, the length step relative to a hs.

### Adequacy between experimental points and theoretical layout

The adjustment is done visually by the trial and error method by searching for the values of the determination coefficient (r^2^) closest to 1 and allocating to the model data values compatible with those in the literature and those used in the calculations of the other 5 accompanying Papers.

## Statistics

See Methods section of Papers 1 and 4.

## Results

### Tension rise

With the values exposed to the first purple column of Table 1, the instantaneous rise of the tension (T) up to the isometric tetanus plateau (T0) is calculated according to (1). The time relationship is represented in Fig 1 by a continuous line for each of the 9 length steps from −1.5 nm to −13.5 nm with a pitch of −1.5 nm. On each of the 9 graphs appear about ten points of the same color as the solid line; the points come from the corresponding curves in Fig 11 from [2]. A good fit between the experimental and theoretical points is observed (r^2^> 99.5%) with the exception of the −1.5 nm step (r^2^=97.9%).

The beginning of the rise is analyzed over the first milliseconds for two types of fibers extracted from the *tibialis anterior* muscle in two species of frogs (Figs 2 and 3 with values taken from the blue and green columns of Table 1). A correct adjustment between experimental and theoretical tensions (r^2^> 97.5%) is obtained, the points distinguished on Figs 2 and 3 coming from the original plots in Fig 12 from [2] and in Fig 2 from [8], respectively.

### End of Phase 2

There is no exact definition of the end of phase 2. We follow the instructions provided by L.E. Ford and coauthors on page 471 in [2]. The tension observed at the end of phase 2 (T2) is equal to the maximum value in the first 15 milliseconds; in the absence of this maximum, an inflection point is sought and T2 corresponds to the intersection of the curve with the tangent to the inflection point.

The stroke size, i.e. the maximum step of a myosin head during the WS, (δX_Max_) varies between 11 and 12 nm (Table 1). A computer routine is designed to find the maximum tension value in the first 15 milliseconds for a length step per hs (ΔX) ranging from 0 to –δX_Max_ with an increment of −0.5 nm. If a maximum is not detected, an inflection point is searched and the previous tension value is taken for T2. The theoretical plot of pT2 (T2/T0) is represented by a continuous red line in Figs 4a and 4b; the data used in the computer routine correspond belong to columns 1 and 3 of Table 1, respectively. The theoretical curve is divided into several parts common to both plots:

**Fig 4.**
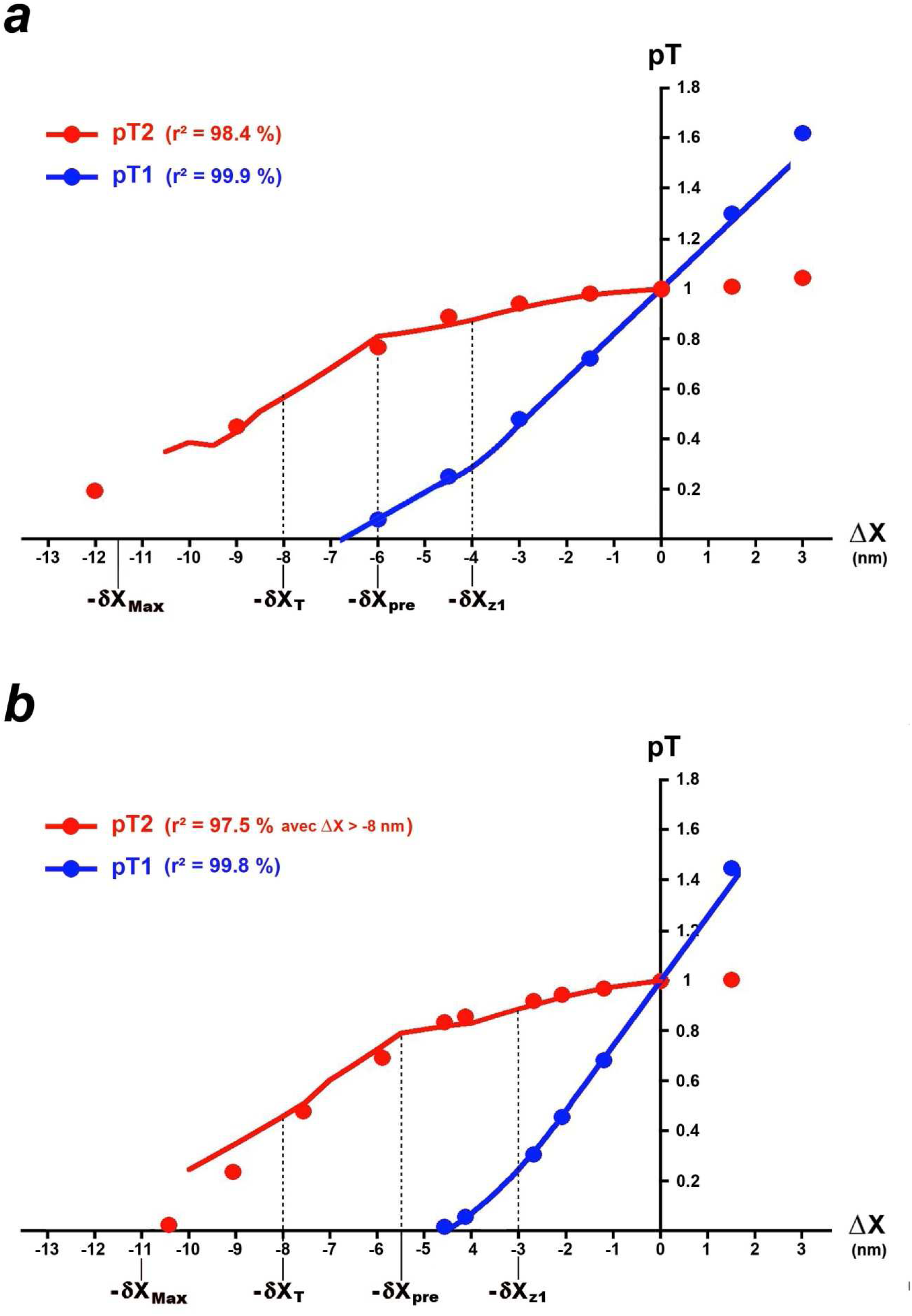
Theoretical tensions determined at the end of phase 1 (blue solid line) and phase 2 (red solid line) as a function of the step size per hs (ΔX). (a) Calculations based on data from a fiber isolated from the *tibialis anterior* muscle of *rana Temporaria* (blue column in Table 1). The points come from Fig 13 in [2]. (b) Calculations based on data from a fiber isolated from the *tibialis anterior* muscle of *Rana Esculenta* (green column in Table 1). The points are from Fig 3B in [4].

1/ −4 nm ≤ ΔX ≤ 0 (parabolic arc)

The end of phase 2 (τ_p2_) occurs for τ_p2_ ≈ 4·τ_startF_ (1-e^-4^=98%), i.e. with τ_startF_ ≈ 0.7 ms (Table 1):

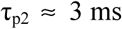

Between 0 and –δX_z1_, the tension is the sum of two linear equations, one of the 1st order with T1_Elas_ formulated in (K17a) and the other of the 2nd order with pT1_d1_ given in (K7). By setting the time at the end of phase 2 (t=τ_p2_), pT2 is equal to:

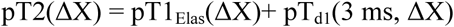

This equation corresponds to a parabola that is extended up to the abscissa equal to −4 nm.

2/ -δX_pre_ ≤ ΔX ≤ −4 nm; approximately straight line segment with a low slope.

3/ -δX_T_ ≤ ΔX ≤ -δX_pre_; approximately straight line segment with a steep slope.

The abscissa -δX_pre_ indicates a slope break: the instantaneous number of heads likely to initiate quickly is maximum relative to the range δX_pre_ (Fig K1b’). Consequently, the tension generated by the new heads decreases sharply for any step ΔX lower than -δX_pre_ (Fig K1c’); the tension increases temporally but the rise is less and less marked (Figs 1, 2 and 3).

4/ −10 nm ≤ ΔX ≤ -δX_T_; extension of the previous segment with a slightly lower slope (Fig 4b) or zigzag (Fig 4a).

5/ ΔX ≤ −10 nm; the algorithm does not detect any maximum or inflection point.

The points shown in Figs 4a and 4b are from Fig 13 in [2] and from Fig 3B in [4], respectively. A correct fit (r^2^ > 97%) is noted for the steps checking ΔX ≥ −8 nm or ΔX ≥ -δX_T_. Below −8nm, there is a discrepancy between the theoretical curve and the experimental dots measured with the naked eye (Fig 4b).

## Discussion

Equation (1), which models the tension kinetics after the end of phase 1, is the sum of various contributions, each associated with a particular event. Some occurrences are concomitant, others follow one another over time. It becomes possible to interpret the 3 phases that classically contribute to the rise from T1 to T0.

### Phase 2

In our model, the rapid rise characteristic of phase 2 has two origins: 1/ the relaxation phenomenon caused by the cancellation of viscosity forces, 2/ the rapid initiation of new myosin heads towards the WS state with the event {startF}.

#### End of phase 2

The end of phase 2 occurs at the end of these two occurrences, on which the beginning of the detachment in fast mode is superimposed with the event {FastDE}. The relative tension at the end of phase 2 (pT2) is algorithmically searched from equation (1) over the first 15 milliseconds of the rise. The theoretical plots in Figs 4a and 4b calculated using the data in Table 1 fit correctly with the experimental points for steps greater than −8 nm (r^2^ > 97%); a divergence appears below −8 nm where our model overestimates the pT2 value (Fig 4b). The model has the classic look of the representative pT2 curve with, on the one hand, a parabolic arc between 0 and −4 nm, and on the other hand, a slope break at the abscissa -δX_pre_ (Figs 4a and 4b).

#### Strong Binding event

The Strong Binding event {SB} is presented in Paragraph B.4 of Supplement S1.B to Paper 1, this event is introduced as a preparatory step for the rapid initiation of a WS with the {startF} event. Before the length step, the myosin heads likely to initiate a WS quickly are first strongly bound according to {SB} but are geometrically unable to achieve a WS because the θ angle of their levers would be geometrically beyond θ_up_, i.e. outside the domain of WS occurrence defined by the interval [θ_down_; θ_up_]. After a consecutive shortening to a length step, heads quickly initiate a WS according to {startF} and the θ angle of their levers is distributed uniformly over an angular range included in the the interval [θ_down_; θ_up_]; see Figs K2b to K2g of Supplement S5.K. The maximum length of this range is δθ_pre_ (Fig K2a). By affine transformation, δθ_pre_ corresponds to the linear range δX_pre_. A slope break is observed between −5 and −6 nm in published curves for pT2; Fig 7 from [8] and Figs 3C and 3D from [9] in frog; Fig 3 from [10] in rat. Our model provides an explanation for the slope change characteristic of the pT2 curve and delivers the value of the abscissa -δX_pre_ equal to the length step per hs corresponding to this break.

With the relationship (K5), the total number of heads likely to initiate a WS according to the 3 modes on the δX_pre_ range (Λ_pre_) is equal to:

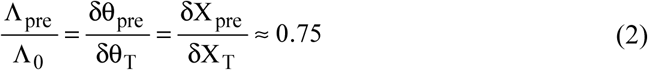

where Λ_0_ is the number of WS myosin heads per hs during the isometric tetanus plateau; δX_T_ and δX_pre_ are two linear ranges equal to 8 and 6 nm, respectively, in accordance with the values presented in tables of Paper 1 and Table 1 in the Results section.

In the same tables are displayed values of p_startF_, the maximum percentage of {startF} realisation, data obtained empirically. The mean value of 60% is chosen:

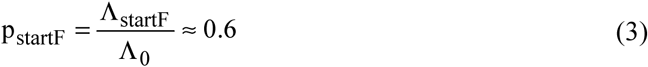

where Λ_startF_ is the maximum number of WS myosin heads per hs that initiated rapidly during the rise to the isometric tetanus plateau.

We deduce from this the maximum percentage (p_SB→startF_) of heads strongly bound during the isometric tetanus plateau and able to quickly initiate a WS:

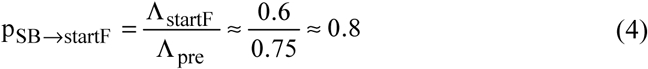

During the isometric tetanus plateau, about 80% of the heads are strongly bound in preparation for a rapid initiation of a WS. Geometrically, we assume that the rapid initiation consists of a succession of Brownian movements of the lever combined with random rotations of this in the OYZ plane perpendicular to OX, the longitudinal axis of the actin filament; when the angle β characteristic of the fixed plane in which the lever of a WS head moves is found, the lever is mechanically guided until it reaches the angular position θ determining the appropriate configuration for the cross-bridge.

#### Independence of pT2 (T2/T0) from T0

If the experimental conditions do not influence the angular ranges, δθ_T_, δθ_pre_ and δθ_Max_, and after affine transformation, the linear ranges, δX_T_, δX_pre_ and δX_Max_, the percentages of equations (2), (3) and (4) are constants. According to the equations in Supplement S5.K, where only percentages, geometric and temporal data are present and constant, the kinetics of the tension rise and in particular the end-of-phase 2 tension referred to T0 (pT2) is not be influenced by T0. This is verified experimentally; see Fig 7 from [11] where T0 varies is a function of the rise duration (300 and 500 ms); superposition of pT2 values for two initial lengths of the sarcomere in Fig 11 from [11]; idem with Fig 3D from [9] where T0 is modified with or without a cross-bridge inhibitor.

#### Independence of pT2 from the duration of phase 1 (τ_p1_)

Section J.9 of the Supplement S4.J to Paper 4 discusses the duration of phase 1 (τ_p1_) as a factor influencing viscosity forces and thus the calculation of T1. Once the effects of viscosity have disappeared during phase 2, T1 becomes T1_Elas_ again, whose value is determined solely on the basis of mechanical linking actions (see Supplement S4.I), a value totally independent of τ_p1_. In this case, pT2 (or T2) depends solely on the rapid initiation of new heads in the blank areas, an event that depends exclusively on the constants p_startF_ and τ_startF_, data also independent of τ_p1_. Therefore pT2 should be independent of τ_p1_, fact consistent with the observations; see Fig 19 in [2].

### Phase 3

The deceleration of tension kinetics, a phenomenon that distinguishes phase 3, comes from several sources: (a) the end of the {startF} event which is expected from time t = 4·τ_startF_ where {startF} reaches 98% of achievements, i.e. 2 to 3 ms after the end of phase 1, (b) the occurrence of the {FastDE} event which occurs as soon as t ≥ τ_preFDE_, i.e. 1 ms after the end of phase 1, an event negatively involved in the tension assessment, (c) the appearance of the {SlowDE} event which occurs if t ≥ τ_preSDE_, i.e. 5 to 7 ms after the end of phase 1, an event whose contribution is also negative.

These three occurrences are associated with the slowing down of the tension increase observed on the plots in Figs 1, 2 and 3. The effects of {startF}, {FastDE} and {SlowDE} vary according to the domain and sector concerned and explain the more or less pronounced presence of a plateau or inflection on the curve.

### Phase 4

The slow rise in the characteristic tension of phase 4 results from: (a) the slow initiation of new heads to the WS state with the event {startS} if t > τ_preS_, (b) the very slow initiation of new heads into the WS state with the event {startVS} if t > τ_preVS_, (c) the stopping of the event {FastDE} which appears from time t = 4·τ_FDE_ where {FastDE} reaches 98% of realisations, i.e. 12 to 20 ms after the end of phase 1, (d) the progressive cessation of the event {SlowDE} from time t = 4·τ_SDE_ where {SlowDE} reaches 98% of achievements, i.e. 40 to 80 ms after the end of phase 1.

For the 9 examples in Fig 1, the tension tends towards T0 after 180 ms according to what is observed in the literature and the explanation provided by the model at the end of the 6 sub-paragraphs K.5.1 to K.5.6 for the 6 sectors studied. After a shortening of the fiber, the spatial distribution of the angular positions θ of the levers belonging to the WS heads in a hs converges temporally towards a uniform density over the interval δθ_T_, a law characteristic of stable equilibrium during the isometric tetanus plateau (see accompanying Papers 3 and 4).

### Large length steps

For large length steps, i.e. ΔX ≤ -(δX_Max_+δX_pre_), a slack of the fiber is observed at the end of phase 1. This phenomenon is interpreted at the end of the Discussion section of Paper 4. Once the effects of viscosity have disappeared, the tension rise is carried out according to the proposed scheme with equations (K11) and (K27) independent of the ΔX level in Domain 4 and Sector 6. The redevelopment of T is close to that given as an example in Fig 1 for the −13.5 nm step belonging to Sector 4.

## Conclusion

On the basis of classical events contributing to the cross-bridge cycle and supporting geometric and temporal data specific to a vertebrate myosin II head, the complete evolution of the tension measured at the end of a muscle fiber is modelled for any length step that verifies “ΔL< 0”. The presence of viscosity during phase 1 and its abolition during phase 2 allows this result.

## Supporting information

Supplementary Chapter. Calculations relating to the increase in tension T to T0 after phase 1 of a length step.

Computer Programs relative to the calculations and plots in Paper 5.

Data relating to computer programs used for Paper 5.

## Acknowledgements

I thank Professor V. Lombardi for allowing me to reproduce the points presented in Fig 3. I thank Professor G. Piazzesi for allowing me to reproduce the points presented in Fig 4b.

## Supporting information

### S5.K Supplementary Chapter. Calculations relating to the increase in tension T to T0 after phase 1 of a length step

K.1 Recalls of the results established in Supplement S1.B of Paper 1

K.2 Calculations of the tension differential created by heads initiating a WS

K.3 Calculations of the tension differential created by rapidly detaching heads

K.4 Calculations of the tension differential created by slowly detaching heads

K5 Establishing the master equation of the model

References of Supplement S5.K

### CP5 Supplementary Material. Computer Programs relative to the calculations and plots in Paper 5

Algorithms are written in Visual Basic 6.

### DA5 Supplementary Material. Data relating to computer programs used for Paper 5

Access Tables are transferred to Excel sheets.

