## Supplementary Chapter. Calculations relating to the increase in tension T to T0 after phase 1 of a length step. for "Mechanical model of muscle contraction. 5. Tension rise after phase 1 of a length step"

### S5.K Supplementary Chapter of Paper 5. Theoretical rise to T0

***For the record, the angle $\theta$ indicates the position of the lever of a head in working stroke (WS) between the two terminals $\theta_{up}$ and $\theta_{down}$ relative to the up and down positions.***

#### K.1 Recalls of the results established in Supplement S1.B of Paper 1

##### *K.1.1 Initiation of the WS state*

The generic WS initiation event {WSstart} is available in 3 modes, {startF}, {startS} or {startVS}, which refer to a Fast, Slow or Very Slow initiation, respectively. There are two cases depending on whether the {startF} event is achievable or not.

**Case 1 where the occurrence of {startF} is possible**

The instantaneous probability of {WSstart} realisation is formulated in Case 1:

$$P_{WS1}(t) = p_{startF} \cdot \left(1 - e^{-\frac{t}{\tau_{startF}}}\right) + p_{startS} \cdot \left(1 - e^{-\frac{t-\tau_{preS}}{\tau_{startS}}}\right) + \left(1 - p_{startF} - p_{startS}\right) \cdot \left(1 - e^{-\frac{t-\tau_{preVS}}{\tau_{startVS}}}\right) \quad \text{(K1a)}$$

where $p_{startF}$ and $p_{startS}$ are the probabilities of occurence of the events {startF} and {startS} after an infinite time, i.e. after several ten milliseconds required by the advent of the isometric tetanus plateau; $\tau_{startF}$, $\tau_{startS}$ and $\tau_{startVS}$ are the time constants for {startF}, {startS} and {startVS}, respectively; $\tau_{preS}$ and $\tau_{preVS}$ are the occurrence delays of {startS} and {startVS}.

**Case 2 where the occurrence of {startF} is impossible**

The instantaneous probability of achievement of {WSstart} is formulated in Case 2:

$$P_{WS2}(t) = \left(p_{startF} + p_{startS}\right) \cdot \left(1 - e^{-\frac{t-\tau_{preS}}{\tau_{startS}}}\right) + \left(1 - p_{startF} - p_{startS}\right) \cdot \left(1 - e^{-\frac{t-\tau_{preVS}}{\tau_{startVS}}}\right) \quad \text{(K1b)}$$

Note that $P_{WS1}$ and $P_{WS2}$ tend towards 1 if t tends towards $+\infty$.

##### *K.1.2 Detachment in Fast mode*

The fast detachment event {FastDE} concerns WS heads whose lever has an angular position $\theta$ beyond $\theta_{down}$ after a length step. In our model, the instantaneous probability of {FastDE} realisation is only formulated in Zone 2 between $-\delta X_{Max}$ and $-\delta X_{z1}$ as:

$$P_{FDE}(t) = \left(1 - e^{-\frac{t-\tau_{preFDE}}{\tau_{FDE}}}\right) . \mathbb{1}_{[-\delta X_{Max};-\delta X_{z1}]}(\Delta X) \quad \text{(K2)}$$

where $\tau_{FDE}$ and $\tau_{preFDE}$ are the time constant and occurrence delay of {FastDE}; $\delta X_{Max}$ is the stroke size; $\delta X_{z1}$ is a linear ranges defined in paragraph I.6 of Supplement S4.I to Paper 4.

##### *K.1.3 Detachment in Slow mode*

During the isometric tetanus plateau, the spatial density of θ is uniformly distributed over an angular range ($\delta\theta_T$) equal to about 50° and framed by the angles $\theta_{up}$ and $\theta_T$ (see Paper 4). The slow detachment event {SlowDE} only concerns WS heads whose lever has an angular position θ varying between $\theta_T$ and $\theta_{down}$.

There are several cases.

**Option 1 where the WS state is present before the length step**

For this option, the instantaneous probability of occurrence of {SlowDE} is called as $P_{SDE\_T}$ (T for Tetanus) and is formulated:

$$P_{SDE\_T}(t) = \left(1 - e^{-\frac{t-\tau_{preSDE}}{\tau_{SDE}}}\right) \quad \text{(K3a)}$$

where $\tau_{SDE}$ and $\tau_{preSDE}$ are the time constant and occurrence delay of {SlowDE}.

**Option 2 with Case 1 where the occurrence of {startF} is possible after the length step**

For this option, the instantaneous probability of achievement of {SlowDE} is named as $P_{SDE\_1}$ and is formulated:

$$P_{SDE\_1}(t) = p_{startF} \cdot \left(1 - e^{-\frac{t-\tau_{preSDE}}{\tau_{SDE}}}\right) + p_{startS} \cdot \left(1 - \frac{\tau_{startS} \cdot e^{-\frac{t-(\tau_{preS}+\tau_{preSDE})}{\tau_{startS}}}}{(\tau_{startS} - \tau_{SDE})} - \frac{\tau_{SDE} \cdot e^{-\frac{t-(\tau_{preS}+\tau_{preSDE})}{\tau_{SDE}}}}{(\tau_{SDE} - \tau_{startS})}\right)$$
$$+ (1 - p_{startF} - p_{startS}) \cdot \left(1 - \frac{\tau_{startVS} \cdot e^{-\frac{t-(\tau_{preVS}+\tau_{preSDE})}{\tau_{startVS}}}}{(\tau_{startVS} - \tau_{SDE})} - \frac{\tau_{SDE} \cdot e^{-\frac{t-(\tau_{preVS}+\tau_{preSDE})}{\tau_{SDE}}}}{(\tau_{SDE} - \tau_{startVS})}\right) \quad \text{(K3b)}$$

**Option 2 with Case 2 where the occurrence of {startF} is impossible after the length step**

For this option, the instantaneous probability of achievement of {SlowDE} is called as $P_{SDE\_2}$ and is formulated:

$$P_{SDE\_2}(t) = (p_{startF} + p_{startS}) \cdot \left(1 - \frac{\tau_{startS} \cdot e^{-\frac{t-(\tau_{preS}+\tau_{preSDE})}{\tau_{startS}}}}{(\tau_{startS} - \tau_{SDE})} - \frac{\tau_{SDE} \cdot e^{-\frac{t-(\tau_{preS}+\tau_{preSDE})}{\tau_{SDE}}}}{(\tau_{SDE} - \tau_{startS})}\right)$$
$$+ (1 - p_{startF} - p_{startS}) \cdot \left(1 - \frac{\tau_{startVS} \cdot e^{-\frac{t-(\tau_{preVS}+\tau_{preSDE})}{\tau_{startVS}}}}{(\tau_{startVS} - \tau_{SDE})} - \frac{\tau_{SDE} \cdot e^{-\frac{t-(\tau_{preVS}+\tau_{preSDE})}{\tau_{SDE}}}}{(\tau_{SDE} - \tau_{startVS})}\right) \quad \text{(K3c)}$$

Note that $P_{SDE\_T}$, $P_{SDE\_1}$ and $P_{SDE\_2}$ tend towards 1 if t tends towards +∞.

#### K. 2 Calculations of the tension created by heads initiating a WS

***General rule for calculating the tension generated by myosin heads initiating a WS in the areas freed after a length step***

Following a disturbance by a length step, the study of phase 1 in Paper 4 shows that each half-sarcomere (hs) is shortened by a variable length due to the presence of viscosity, a length that remains close to the mean value ($\overline{\Delta X}$). After phase 1, the effects of viscosity disappear quickly (< 0.5 ms) and at this point all hs of the fiber are in isometry. The only actions then present are the linking forces and moments; this case is studied in paragraph I.5 of Supplement S4.I to Paper 4. It appears that at any moment, each hs of the fiber has an equal number of WS heads, an identical distribution of the angle θ, an equality of the tensions ($T_{hs}$) acting at the two edges of the hs alternately delimited by a Z-disk and a M-disk, and a shortening (ΔX) common to all hs such as:

$$\Delta X = \overline{\Delta X} = \frac{\Lambda L}{N_{hs}}$$

where ΔL is the shortening of the fiber; $N_{hs}$ is the number of hs per myofibril.

The results of accompanying Paper 3 are applicable, in particular the homogeneity corollary of hypothesis 5 which stipulates that the distribution of θ within each hs follows exactly the same uniform law whatever the length of the hs when this length is between 1 and 1.05 µm, and consequently after a step lower than about ten nanometers in modulus. The hs shortening (ΔX) causes the rotation (Δθ) of the levers depending of myosin heads previously in WS and frees up blank areas (Fig K1), i.e. free of WS heads. In these blank intervals included in the maximal angular range ($\delta\theta_{Max}$), heads initiate a WS according to one of the 3 modes, {startF}, {startS} or {startVS}, discussed in paragraph K.1. By passing from discrete to continuous, the angle θ of the levers belonging to these heads is associated to Θ, the continuous random variable distributed uniformly over the blank interval $\delta\theta_L$ according to the law ($\mathcal{U}_L$) defined in (15) in Paper 3:

$$\mathcal{U}_L(\theta) = \frac{1}{\delta\theta_L}.\mathbf{1}_{[\theta_2;\theta_1]}(\theta) \qquad \text{(K4)}$$

where $\theta_1$ and $\theta_2$ are two angles that check the condition " $\theta_{down} \le \theta_2 < \theta_1 \le \theta_{up}$ " in a hs on the right and the condition " $\theta_T \le \theta_1 \le \theta_2 \le \theta_{down}$ " in a hs on the left; $\delta\theta_L$ is an angular range equal to $|\theta_2 - \theta_1|$.

The two angles $\theta_1$ and $\theta_2$ will be determined for each of the intervals or domains studied.

At an infinite time, i.e. several dozen milliseconds in reality, the total number of heads ($\Lambda_{WSstart}$) who have initiated a WS in the range $\delta\theta_L$ between $\theta_{up}$ and $\theta_{down}$ verifies the homogeneity relationship given in (I11) in supplement S4.I of Paper 4 according to:

$$\Lambda_{WSstart} = \Lambda_0 \cdot \frac{\delta\theta_L}{\delta\theta_T} \qquad \text{(K5)}$$

where $\Lambda_0$ is the number of WS myosin heads per hs during the isometric tetanus plateau.

**If Case 1 in sub-paragraph K.1.1 is checked with the possibility of {startF} occuring**

The instantaneous number of heads initiating a WS ($\Lambda_{WS1}$) is equal to:

$$\Lambda_{WS1}(t) = P_{WS1}(t) \cdot \Lambda_{WSstart}$$

where $P_{WS1}$ is a probability determined in (K1a).

In support of the theoretical developments leading to equation (I31b) in paragraph I.4 of Supplement S4.I, the contribution of $\Lambda_{WS1}$ heads to the instantaneous relative tension ($pT_{WS1}$) is formulated with (K5) and the previous equality:

$$pT_{WS1}(t) = P_{WS1}(t) \cdot \left(\frac{\delta X_L}{\delta X_T}\right) \cdot \left[1 + \frac{(X_1 + X_2)}{2 \cdot |X_{down}|}\right] \tag{K6a}$$

where $X_1$ and $X_2$ are the two abscissa corresponding to the two angles $\theta_1$ and $\theta_2$ defined in (K4) according to the affine relation provided in (I21a) in Supplement S4.I.

**If Case 2 in sub-paragraph K.1.1 is checked with the impossibility of {startF} occuring**

The instantaneous number of heads initiating a WS ($\Lambda_{WS2}$) is equal to:

$$\Lambda_{WS2}(t) = P_{WS2}(t) \cdot \Lambda_{WSstart}$$

where $P_{WS2}$ is a probability determined in (K1b).

The contribution to the instantaneous relative tension ($pT_{WS2}$) is formulated:

:

$$pT_{WS2}(t) = P_{WS2}(t) \cdot \left(\frac{\delta X_L}{\delta X_T}\right) \cdot \left[1 + \frac{(X_1 + X_2)}{2 \cdot |X_{down}|}\right] \tag{K6b}$$

We are interested in the domains freed by the heads previously in WS during the isometric tetanus plateau preceding the hs shortening following a length step. The two intervals involved to characterize the domains are $\delta X_{pre}$ and $\delta X_{Max}$: $\delta X_{pre}$ is a linear range defined in the Methods section of accompanying Paper 1 and $\delta X_{Max}$ is the stroke size or maximum step of a head during the WS.

**Reminder:** if the strongly linked heads likely to initiate a WS quickly were in WS before the length step, then the $\theta$ angle of their levers would have a value higher (lower) than $\theta_{up}$ in a hs on the right (left) and would be distributed uniformly in the interval $[\theta_{up} ; (\theta_{up} + \delta\theta_{pre})]$ ( $[(\theta_{up} - \delta\theta_{pre}) ; \theta_{up}]$).

In the linear domain bounded by $\theta_{up}$ and $\theta_{down}$ (Fig 1a), the range $\delta\theta_{pre}$ is linearly transformed into $\delta X_{pre}$ (Fig 1a') according to the relationship (9) of Paper 4.

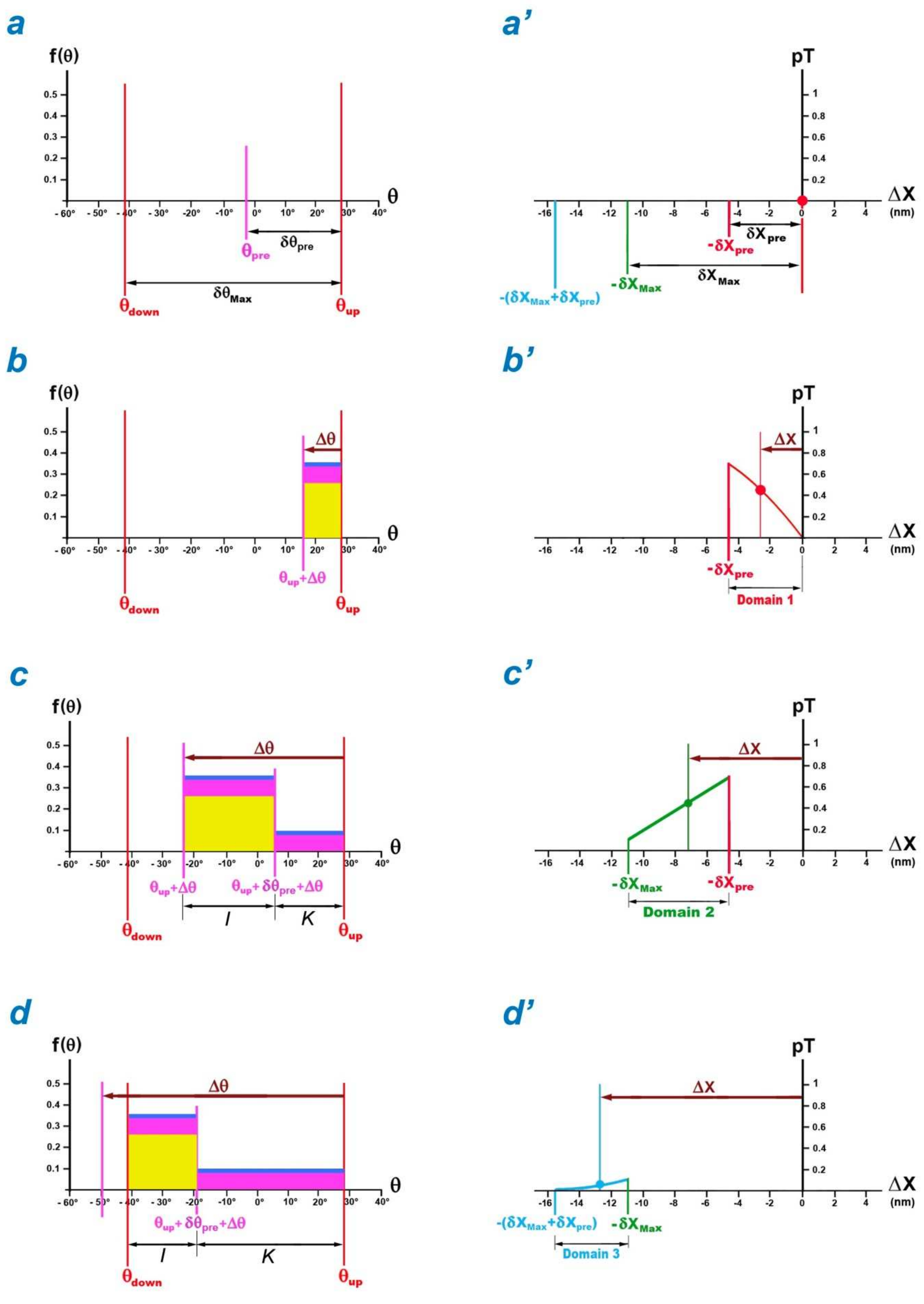

**Fig K1 : Tension created by heads initiating a WS in a half-sarcomere on the right.**

(a) Angular ranges $\delta\theta_{pre}$ and $\delta\theta_{Max}$. (a') Isometric tetanus plateau ($\Delta X=0$) with the 2 linear ranges $\delta X_{pre}$ and $\delta X_{Max}$. (b), (c) and (d) Uniform densities of $\theta$ at time t=3 ms after lever rotation ($\Delta\theta$). (b'), (c') and (d') Relative tension as a function of the length step per hs ($\Delta X$) in Domains 1, 2 and 3, respectively, at time t=3 ms.

##### *K.2.1 Domain 1 ≡ [ $-\delta X_{pre}$ ; 0 [*

The hs shortening ($\Delta X$) is framed (Fig K1b'):

$$-\delta X_{pre} \le \Delta X \le 0$$

By affine transformation, the rotation ($\Delta\theta$) of the levers belonging to heads previously in WS verifies the corresponding inequalities:

$$-\delta\theta_{pre} \le \Delta\theta \le 0$$

In the angular range released and bounded by ($\theta_{up}+\Delta\theta$) and $\theta_{up}$, the $\theta$ angle of the levers belonging to the heads initiating a WS is distributed uniformly on $|\Delta\theta|$. The corresponding Uniform law according to (K4) is represented in proportion on Fig K2b by a yellow rectangle for the fast mode {startF}, a purple rectangle for the slow mode {startS}, and a flattened blue rectangle for the very slow mode {startVS}. After the length step, the occurrence of {startF} is possible: this is Case 1 for which the instantaneous probability of {WSstart} realisation is determined in (K1a).

**Conditions:**

$$\left.\begin{array}{r} p = P_{WS1}(t) \\ X_1 = X_{up} \\ X_2 = X_{up} + \Delta X \end{array}\right\} \Rightarrow \begin{array}{c} \delta X_L = |\Delta X| \\ \\ X_1 + X_2 = 2X_{up} + 2\Delta X \end{array}$$

**Calculation of the tension in range 1 ($pT_{d1}$) with application of (K6a):**

$$pT_{d1}(t,\Delta X) = P_{WS1}(t) \cdot \frac{|\Delta X| \cdot \left[\delta X_{Max} + \frac{\Delta X}{2}\right]}{|X_{down}| \cdot \delta X_T} \quad \text{(K7)}$$

At any time t in Domain 1, the relationship between $pT_{d1}$ and $\Delta X$ is represented by a concave parabolic arc (Fig K1b'; red line).

##### *K.2.2 Domain 2 ≡ [ $-\delta X_{Max}$ ; $-\delta X_{pre}$ [*

The hs shortening ($\Delta X$) verifies the inequalities:

$$-\delta X_{Max} \le \Delta X < -\delta X_{pre}$$

By affine transformation, the rotation ($\Delta\theta$) of levers belonging of the head previously in WS corresponds as:

$$-\delta\theta_{Max} \le \Delta\theta < -\delta\theta_{pre}$$

The pivoting of the levers releases an angular range which is divided into 2 intervals named as I and K (Fig. K1c).

1/ An interval I of $\delta\theta_{pre}$ range bounded by ($\theta_{up}$+$\Delta\theta$) and ($\theta_{up}$+$\Delta\theta$+$\delta\theta_{pre}$) where heads initiate according to the 3 modes with Case 1.

**Conditions with (K1a):**

$$\left.\begin{array}{r} p = P_{WS1}(t) \\ X_1 = X_{up} + \delta X_{pre} + \Delta X \\ X_2 = X_{up} + \Delta X \end{array}\right\} \Rightarrow \begin{array}{c} \delta X_L = \delta X_{pre} \\ \\ X_1 + X_2 = 2X_{up} + \delta X_{pre} + 2\Delta X \end{array}$$

**Calculation of the tension in the interval I of Domain 2 ($pT_{d2\_I}$) with application of (K6a):**

$$pT_{d2\_I}(t,\Delta X) = P_{WS1}(t) \cdot \frac{\delta X_{pre}}{|X_{down}| \cdot \delta X_T} \cdot \left[\delta X_{Max} + \frac{\delta X_{pre}}{2} + \Delta X\right]$$

2/ A complementary range K interval (|$\Delta\theta$| - $\delta\theta$) between ($\theta_{up}$+ $\Delta\theta$ + $\delta\theta_{pre}$) and $\theta_{up}$ where no head is able to quickly initiate a WS. After the step, the occurrence of {startF} is impossible and Case 2 where the heads initiate only according to {startS} or {startVS} is retained. The probability of {WSstart} realisation is delivered in (K1b).

**Conditions:**

$$\left.\begin{array}{r} p = P_{WS2}(t) \\ X_1 = X_{up} \\ X_2 = X_{up} + \delta X_{pre} + \Delta X \end{array}\right\} \Rightarrow \begin{array}{c} \delta X_L = |\Delta X| - \delta X_{pre} \\ \\ X_1 + X_2 = 2X_{up} + \delta X_{pre} + \Delta X \end{array}$$

**Calculation of the tension in the interval K of Domain 2 ($pT_{d2\_K}$) with application of (K6b):**

$$pT_{d2\_K}(t,\Delta X) = P_{WS2}(t) \cdot \frac{\left(|\Delta X| - \delta X_{pre}\right)}{|X_{down}| \cdot \delta X_T} \cdot \left[\delta X_{Max} + \frac{\delta X_{pre}}{2} + \frac{\Delta X}{2}\right]$$

3/ The two contributions on intervals I and K are added together:

$$pT_{d2}(t,\Delta X) = pT_{d2\_I}(t,\Delta X) + pT_{d2\_K}(t,\Delta X)$$

So:

$$pT_{d2}(t,\Delta X) = P_{WS1}(t) \cdot \frac{\delta X_{pre}}{|X_{down}| \cdot \delta X_T} \cdot \left[\delta X_{Max} + \frac{\delta X_{pre}}{2} + \Delta X\right] + P_{WS2}(t) \cdot \frac{\left(|\Delta X| - \delta X_{pre}\right)}{|X_{down}| \cdot \delta X_T} \cdot \left[\delta X_{Max} + \frac{\delta X_{pre}}{2} + \frac{\Delta X}{2}\right] \quad \text{(K8)}$$

At any given time t in Domain 2, the relationship between $pT_{d2}$ and $\Delta X$ is represented by an almost linear parabolic arc (Fig K1c'; green line).

##### *K.2.3 Domain 3 ≡ [ -($\delta X_{Max}$+$\delta X_{pre}$) ; - $\delta X_{Max}$ [*

The hs shortening ($\Delta X$) checks for inequalities:

$$(\delta X_{Max} + \delta X_{pre}) \leq \Delta X < -\delta X_{Max}$$

By affine relation, the rotation ($\Delta\theta$) of levers belonging to the head previously in WS validates the following inequality:

$$\Delta\theta < -\delta\theta_{Max}$$

The pivoting releases an angular range which is divided into 2 intervals (Fig K1d).

1/ An interval I of extent ($\delta\theta_{Max}$+$\delta\theta_{pre}$ +$\Delta\theta$) bounded by $\theta_{down}$ and ($\theta_{up}$+$\delta\theta_{pre}$ +$\Delta\theta$) where heads initiate according to the 3 coexisting modes.

**Reminders of relations provided in paragraph I.4 of Supplement S4.I:**

$$\delta X_{Max} = X_{up} - X_{down} = \delta X_T + \delta X_E \quad \text{(K9a)}$$

$$X_{up} = \frac{\delta X_T}{2} \quad \text{(K9b)}$$

$$\left|X_{down}\right| = \delta X_E + \frac{\delta X_T}{2} \quad \text{(K9c)}$$

**Conditions with (K1a), (K9a), (K9b) and (K9c):**

$$\left.\begin{aligned} p &= P_{WS1}(t) \\ X_1 &= X_{up} + \delta X_{pre} + \Delta X \\ X_2 &= X_{down} \end{aligned}\right\} \Rightarrow \quad \begin{aligned} \delta X_L &= \delta X_{Max} + \delta X_{pre} + \Delta X \\ X_1 + X_2 &= -\delta X_E + \delta X_{pre} + \Delta X \end{aligned}$$

**Calculation of the tension in the interval I of Domain 3 ($pT_{d3\_I}$) with application of (K6a), (K9a), (K9b) and (K9c):**

$$pT_{d3\_I}(t,\Delta X) = P_{WS1}(t) \cdot \frac{\left(\delta X_{Max} + \delta X_{pre} + \Delta X\right)^2}{2 \cdot \left|X_{down}\right| \cdot \delta X_T}$$

2/ A complementary interval K of extent (|$\Delta\theta$| - $\delta\theta$) between ($\theta_{up}$+ $\Delta\theta$ + $\delta\theta_{pre}$) and $\theta_{up}$ where no head is able to quickly initiate a WS, case studied previously. In this interval, the calculation of the tension is identical to that performed in interval K of Domain 2 as:

$$pT_{d3\_K}(t,\Delta X) = P_{WS2}(t) \cdot \frac{\left(\left|\Delta X\right| - \delta X_{pre}\right)}{\left|X_{down}\right| \cdot \delta X_T} \cdot \left[\delta X_{Max} + \frac{\delta X_{pre}}{2} + \frac{\Delta X}{2}\right]$$

3/ By summing the 2 contributions on the 2 intervals I and K, we obtain:

$$pT_{d3}(t,\Delta X) = pT_{d3\_I}(t,\Delta X) + pT_{d3\_K}(t,\Delta X)$$

So:

$$pT_{d3}(t,\Delta X) = P_{WS1}(t) \cdot \frac{\left(\delta X_{Max} + \delta X_{pre} + \Delta X\right)^2}{2 \cdot \left|X_{down}\right| \cdot \delta X_T} + P_{WS2}(t) \cdot \frac{\left(\left|\Delta X\right| - \delta X_{pre}\right)}{\left|X_{down}\right| \cdot \delta X_T} \cdot \left[\delta X_{Max} + \frac{\delta X_{pre}}{2} + \frac{\Delta X}{2}\right] \quad \text{(K10)}$$

At any time t in Domain 3, the relationship between $pT_{d3}$ and $\Delta X$ is representad by a convex parabolic arc (Fig K1c'; purple line).

##### *K.2.4 Domain 4 where $\Delta X < -(\delta X_{Max}+\delta X_{pre})$*

The hs shortening ($\Delta X$) verifies:

$$\Delta X < -(\delta X_{Max} + \delta X_{pre})$$

The inequality becomes by affine transformation:

$$\Delta\theta < -\delta\theta_{Max}$$

Fast initiation of a WS with {startF} event is not possible. The heads initiate a WS slowly or very slowly according Case 2 in the interval $\delta\theta_{Max}$ limited by $\theta_{down}$ and $\theta_{up}$.

**Conditions with (K1b), (K9b) and (K9c):**

$$\left.\begin{array}{r} p = P_{WS2}(t) \\ X_1 = X_{up} \\ X_2 = X_{down} \end{array}\right\} \Rightarrow \begin{array}{c} \delta X_L = \delta X_{Max} \\ \\ X_1 + X_2 = -\delta X_E \end{array}$$

**Calculation of the tension in Domain 4 ($pT_{Ritp}$ ; Ritp for *Rise to isometric tetanus plateau*) with application of (K6b), (K9a) and (K9c):**

$$pT_{Ritp}(t) = P_{WS2}(t) \cdot \frac{(\delta X_{Max})^2}{2 \cdot |X_{down}| \cdot \delta X_T} \qquad \text{(K11)}$$

The relationship (K11) is independent of $\Delta X$.

This temporal equation characterizes the tension rise up to the isometric tetanus plateau which follows a significant step after the disappearance of the *slack* phenomenon (see accompanying Paper 4)

#### K. 3 Detachment according to fast mode {FastDE}

In paragraph I.5 of Supplement S4.I to Paper 4, two calculations of the tension at the end of phase 1 in Zone 2 ( $pT1^{P}_{Elas,z2}$ and $pT1^{S}_{Elas,z2}$ ) are carried out: see equations (I47) and (I48) illustrated with Fig I2c' and I2d', respectively. A compromise solution is chosen in paragraph I.6 with equation (I53a). After phase 1, the heads are detaching quickly according to the event {FastDE} and consecutively the tension tends towards $pT1^{P}_{Elas,z2}$, the tension for which all the heads whose levers have an angular position beyond $\theta_{up}$ are detached. The instantaneous contribution caused by the rapid detachment ($\Delta pT_{FDE}$) is determined by making the difference between the tensions formulated in (I53a) and (I47), calculated pro rata to the instantaneous number of heads involved by the event {FastDE}, that is :

$$\Delta pT_{FDE}(t,\Delta X) = P_{FDE}(t) \cdot \left[ \chi_{z2} \cdot (\delta X_{Max} + \Delta X) - \frac{(\delta X_{Max} + \Delta X)^2}{2 \cdot \delta X_T \cdot |X_{down}|} \right] \cdot \mathbf{1}_{[-\delta X_{Max};-\delta X_{z1}]}(\Delta X) \qquad \text{(K12)}$$

where $P_{FDE}$ is the instantaneous probability of {FastDE} achievement given in (K2); $\chi_{z2}$ is the elastic origin slope in Zone 2 formulated with (I53b) and reproduced below:

$$\chi_{z2} = \frac{(1 - \chi_{z1} \cdot \delta X_{z1})}{\delta X_{Max} - \delta X_{z1}}$$

and where $\delta X_{z1}$ is an angular range introduced in paragraph I.6 to redefine Zones 1 and 2 (Fig I3).

#### K. 4 Detachment in slow mode {SlowDE}

This event only concerns WS heads whose lever has an angular position θ between $\theta_T$ and $\theta_{down}$. However, depending on the value of the length step, the WS state of these myosin heads has different origins.

##### *General rule for calculating the tension generated by slowly detaching myosin heads*

All WS heads with a lever angle θ between $\theta_{down}$ and $\theta_T$ detach slowly and uniformly in any $\delta\theta_L$ interval included in $\delta\theta_E$ (Fig K2a). By passing from discrete to continuous, the angle θ of the levers belonging to these heads is associated with Θ, the random variable continues distributed uniformly on $\delta\theta_L$ in accordance with the formulation delivered in (15) in accompanying Paper 3, that is:

$$\mathcal{U}_L(\theta) = \frac{1}{\delta\theta_L} . \mathbf{1}_{[\theta_2;\theta_1]}(\theta) \qquad \text{(K13)}$$

where $\theta_1$ and $\theta_2$ are two angles that verify the condition " $\theta_{down} \le \theta_2 \le \theta_1 \le \theta_T$ " in a hs on the right and the condition " $\theta_T \le \theta_1 \le \theta_2 \le \theta_{down}$ " in a hs on the left; $\delta\theta_L$ is an angular range equal to $|\theta_2 - \theta_1|$.

The two angles $\theta_1$ and $\theta_2$ will be determined for each of the sectors studied.

At an infinite time, i.e. several dozen milliseconds in reality, the total number of heads ($\Lambda_{SDE}$) previously in WS that have slowly detached after the length step in the range $\delta\theta_L$ between $\theta_T$ and $\theta_{down}$ verifies the homogeneity relationship given in (I11) in Supplement S4.I of Paper 4 as:

$$\Lambda_{SDE} = \Lambda_0 \cdot \frac{\delta\theta_L}{\delta\theta_T} \tag{K14}$$

**If Option 1 in sub-paragraph K.1.2 is proven**

The instantaneous number of slowly detaching heads ($\Lambda_{SDE\_T}$) is equal to:

$$\Lambda_{SDE\_T}(t) = P_{SDE\_T}(t) \cdot \Lambda_{SDE}$$

where $P_{SDE\_T}$ is an instantaneous probability determined in (K3a).

In support of the theoretical developments leading to equation (I31b) in paragraph I.4 of Supplement S4.I, the contribution of $\Lambda_{SDE\_T}$ heads to the instantaneous relative tension ($pT_{SDE\_T}$) is formulated with (K14) and the previous equality:

$$pT_{SDE\_T}(t) = P_{SDE\_T}(t) \cdot \left(\frac{\delta X_L}{\delta X_T}\right) \cdot \left[1 + \frac{(X_1 + X_2)}{2 \cdot |X_{down}|}\right] \tag{K15a}$$

where $X_1$ and $X_2$ are the two abscissa corresponding to the two angles $\theta_1$ and $\theta_2$ defined in (K13) according to the relation (I21a) delivered in the Supplement S4.I.

**If Option 2 with Case 1 in sub-paragraph K.1.2 is proven**

The instantaneous number of slowly detaching heads ($\Lambda_{SDE\_1}$) is equal to:

$$\Lambda_{SDE\_1}(t) = P_{SDE\_1}(t) \cdot \Lambda_{SDE}$$

where $P_{SDE\_1}$ is an instantaneous probability determined in (K3b).

The contribution of $\Lambda_{SDE\_1}$ heads to the instantaneous relative tension ($pT_{SDE\_1}$) is formulated:

$$pT_{SDE\_1}(t) = P_{SDE\_1}(t) \cdot \left(\frac{\delta X_L}{\delta X_T}\right) \cdot \left[1 + \frac{(X_1 + X_2)}{2 \cdot |X_{down}|}\right] \tag{K15b}$$

**If Option 2 with Case 2 in sub-paragraph K.1.2 is proven**

The instantaneous number of slowly detaching heads ($\Lambda_{SDE\_2}$) is equal to:

$$\Lambda_{SDE\_2}(t) = P_{SDE\_2}(t) \cdot \Lambda_{SDE}$$

where $P_{SDE\_2}$ is an instantaneous probability determined in (K3c).

The contribution of the $\Lambda_{SDE\_2}$ heads to the instantaneous relative tension ($pT_{SDE\_2}$) is formulated:

$$pT_{SDE\_2}(t) = P_{SDE\_2}(t) \cdot \left(\frac{\delta X_L}{\delta X_T}\right) \cdot \left[1 + \frac{(X_1 + X_2)}{2 \cdot |X_{down}|}\right] \tag{K15c}$$

The 4 ranges presented below are ordered according to their values (see Table 1 of Paper 5):

$$\delta X_E < \delta X_{pre} < \delta X_T < \delta X_{Max}$$

In abscissa, this gives in a half-sarcomere on the right:

$$-(\delta X_{Max} + \delta X_{pre}) < -(\delta X_T + \delta X_{pre}) < -\delta X_{Max} < -\delta X_T < -\delta X_{pre} < -\delta X_E$$

**Calculation principle**: an interval between two consecutive abscissa presented above is converted into a sector. The hs shortening ($\Delta X$) is studied according to the sector and we distinguish the WS heads whose lever has an angular position between $\theta_{down}$ and $\theta_T$. We obtain 7 scenarios for 6 sectors (Fig K2b to K2h). In each case, the relative tension generated by these slowly detaching heads according to the option chosen is calculated using equations (K15a), (K15b) or (K15c). Then at an infinite time, it is verified that the tension generated by the heads remaining in WS between $\theta_T$ and $\theta_{up}$ tends towards T0, the tension of the isometric tetanus plateau.

**Reminders of relationships provided in paragraph I.4 of Supplement S4.I:**

$$\delta X_{Max} = \delta X_T + \delta X_E \quad \text{(K16a)}$$

$$X_{up} = \frac{\delta X_T}{2} \quad \text{(K16b)}$$

$$|X_{down}| = \delta X_E + \frac{\delta X_T}{2} \quad \text{(K16c)}$$

**Reminders of equations in paragraph I.5 of Supplement S4.I with the calculation of the relative tension in the absence of viscosity in Zones 1 and 2:**

$$pT_{z1} = 1 + \frac{\Delta X}{|X_{down}|} \quad \text{(K17a)}$$

$$pT_{z2} = \frac{(\delta X_{Max} + \Delta X)^2}{2 \cdot \delta X_T \cdot |X_{down}|} \quad \text{(K17b)}$$

**Reminders of the calculation of the relative tension in Domains 1 to 4 from (K7) to (K11):**

$$pT_{d1}(t, \Delta X) = P_{WS1}(t) \cdot \frac{|\Delta X| \cdot \left[\delta X_{Max} + \frac{\Delta X}{2}\right]}{|X_{down}| \cdot \delta X_T} \quad \text{(K18a)}$$

$$pT_{d2}(t, \Delta X) = P_{WS1}(t) \cdot \frac{\delta X_{pre}}{|X_{down}| \cdot \delta X_T} \cdot \left[\delta X_{Max} + \frac{\delta X_{pre}}{2} + \Delta X\right] + P_{WS2}(t) \cdot \frac{(|\Delta X| - \delta X_{pre})}{|X_{down}| \cdot \delta X_T} \cdot \left[\delta X_{Max} + \frac{\delta X_{pre}}{2} + \frac{\Delta X}{2}\right]$$

(K18b)

$$pT_{d3}(t, \Delta X) = P_{WS1}(t) \cdot \frac{(\delta X_{Max} + \delta X_{pre} + \Delta X)^2}{2 \cdot |X_{down}| \cdot \delta X_T} + P_{WS2}(t) \cdot \frac{(|\Delta X| - \delta X_{pre})}{|X_{down}| \cdot \delta X_T} \cdot \left[\delta X_{Max} + \frac{\delta X_{pre}}{2} + \frac{\Delta X}{2}\right] \quad \text{(K18c)}$$

$$pT_{Ritp}(t) = P_{WS2}(t) \cdot \frac{(\delta X_{Max})^2}{2 \cdot |X_{down}| \cdot \delta X_T} \quad \text{(K18d)}$$

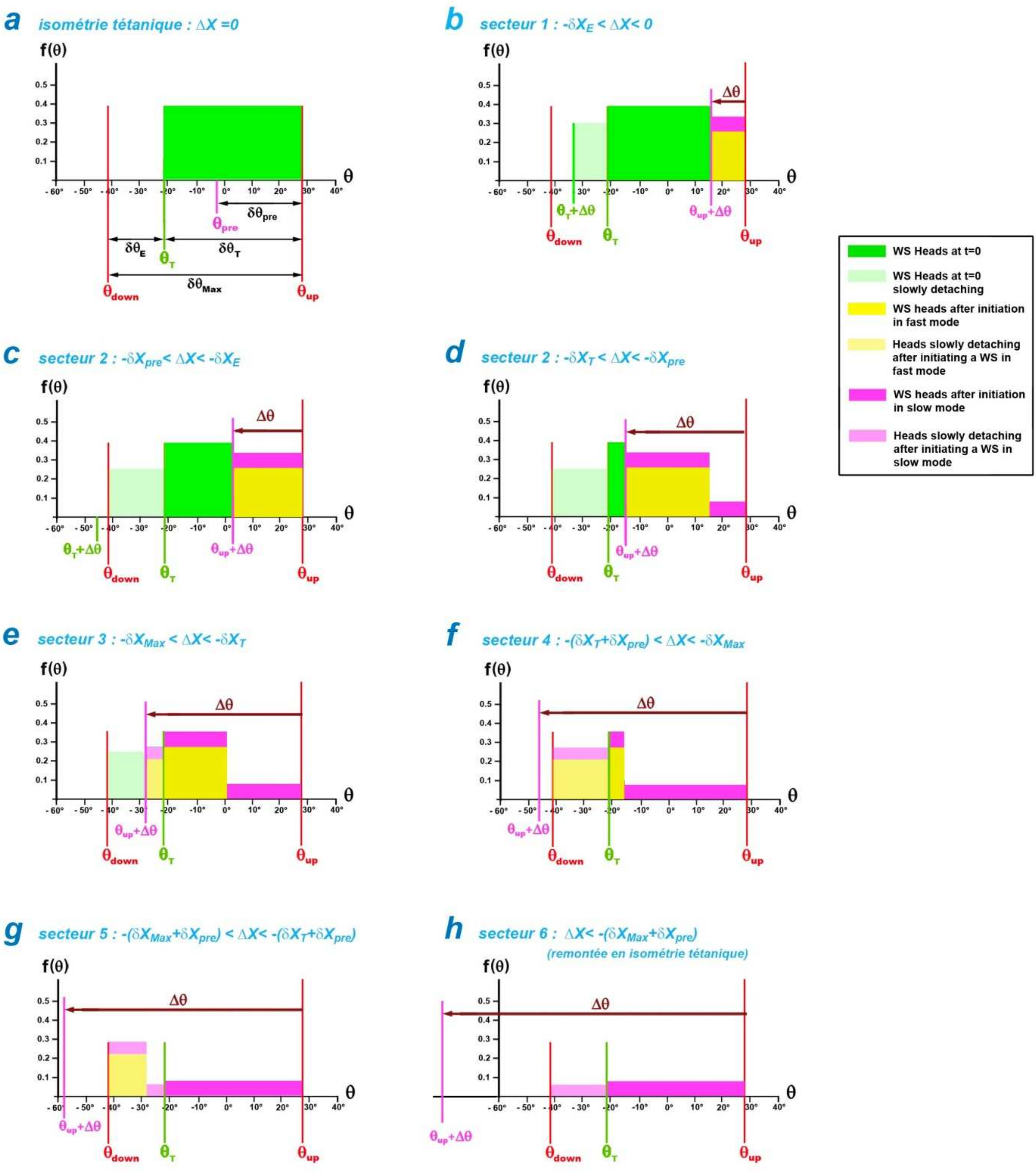


**Fig K2 : Uniform distributions of θ in a right half-sarcomere.**

(a) Isometric tetanus plateau (ΔX=0).

(b), (c), (d), (e), (f), (g) and (h) Uniform densities of θ at time t=15 ms after lever rotation (Δθ) corresponding to the length step per hs (ΔX) in Sectors 1, 2, 2, 3, 4, 5 and 6, respectively.

##### *K.4.1 Sector 1 ≡ [ $-\delta X_E$ ; 0 [*

The hs shortening ($\Delta X$) is framed:

$$-\delta X_E \leq \Delta X < 0$$

By affine transformation, the rotation ($\Delta\theta$) of levers belonging to the heads previously in WS verifies the corresponding inequalities:

$$-\delta \theta_E \leq \Delta\theta < 0$$

All heads with a lever angle $\theta$ between $\theta_{down}$ and $\theta_T$ were in WS state previously at the length step (Fig K2b; light green rectangle).

Option 1 is applied.

**Conditions with (K3):**

$$\left.\begin{array}{r} p = P_{SDE\_T}(t) \\ X_1 = X_T \\ X_2 = X_T + \Delta X \end{array}\right\} \Rightarrow \begin{array}{r} \delta X_L = |\Delta X| \\ \\ X_1 + X_2 = -\delta X_T + \Delta X \end{array}$$

**Calculation of the relative tension in Sector 1 ($pT_{s1}$) with application of (K15a):**

$$pT_{s1}(t, \Delta X) = P_{SDE\_T}(t) \cdot \frac{|\Delta X| \cdot \left[\delta X_E + \frac{\Delta X}{2}\right]}{|X_{down}| \cdot \delta X_T} \qquad \text{(K19)}$$

**<u>Rise up to T0</u>**

Sector 1 coincides with Zone 1 and belongs to Domain 1.

So, for t infinite:

$$pT(t\rightarrow+\infty, \Delta X) = pT_{z1}(\Delta X) + pT_{d1}(t\rightarrow+\infty, \Delta X) - pT_{s1}(t\rightarrow+\infty, \Delta X)$$

With (K17a), (K18a) and (K19):

$$pT(t \rightarrow +\infty, \Delta X) = 1 + \frac{\Delta X}{|X_{down}|} + \frac{|\Delta X| \cdot \left[\delta X_{Max} + \frac{\Delta X}{2}\right]}{|X_{down}| \cdot \delta X_T} - \frac{|\Delta X| \cdot \left[\delta X_E + \frac{\Delta X}{2}\right]}{|X_{down}| \cdot \delta X_T}$$

After reduction and using (K16a):

$$pT(t\rightarrow+\infty, \Delta X) = 1$$

##### *K.4.2 Sector 2 ≡ [ $-\delta X_T$ ; $-\delta X_E$ [*

The hs shortening ($\Delta X$) is framed:

$$-\delta X_T \leq \Delta X < -\delta X_E$$

By affine transformation, the rotation ($\Delta\theta$) of levers belonging to the heads previously in WS verifies the corresponding inequalities:

$$-\delta\theta_T \leq \Delta\theta < -\delta\theta_E$$

All heads with a lever angle $\theta$ between $\theta_{down}$ and $\theta_T$ were previously in WS state (Fig K2b; light green rectangle). Option 1 is still applied.

**Development by squaring the expression (K16a) and with (K16c):**

$$\delta X_{Max}{}^2 - \delta X_E{}^2 = \delta X_T{}^2 + 2 \cdot \delta X_T \cdot \delta X_E = 2 \cdot |X_{down}| \cdot \delta X_T \qquad \text{(K20)}$$

**Conditions using (K3):**

$$\left.\begin{aligned} p &= P_{SDE\_T}(t) \\ X_1 &= X_T \\ X_2 &= X_{down} \end{aligned}\right\} \Rightarrow \quad \begin{aligned} \delta X_L &= \delta X_E \\ X_1 + X_2 &= -\frac{\delta X_T}{2} - |X_{down}| \end{aligned}$$

**Calculation of the relative tension in mains 2 ($pT_{s2}$) with application of (K15a):**

$$pT_{s2}(t, \Delta X) = pT_{s2}(t) = P_{SDE\_T}(t) \cdot \frac{\delta X_E{}^2}{2 \cdot |X_{down}| \cdot \delta X_T} \qquad \text{(K21)}$$

**<u>Rise to T0</u>**

There are two scenarios (Figs K2c and K2d).

**1/ $\Delta X \in$ [ $-\delta X_{pre}$ ; $-\delta X_E$ [**

This part of Sector 2 (Fig K2c) belongs to Zone 2 and Domain 1.

So, for t infinite:

$$pT(t\rightarrow+\infty, \Delta X) = pT_{z2}(\Delta X) + pT_{d1}(t\rightarrow+\infty, \Delta X) - pT_{s2}(t\rightarrow+\infty, \Delta X)$$

With (K17b), (K18a) and (K21):

$$pT(t \rightarrow +\infty, \Delta X) = \frac{(\delta X_{Max} + \Delta X)^2}{2 \cdot \delta X_T \cdot |X_{down}|} + \frac{|\Delta X| \cdot \left[\delta X_{Max} + \frac{\Delta X}{2}\right]}{|X_{down}| \cdot \delta X_T} - \frac{\delta X_E{}^2}{2 \cdot |X_{down}| \cdot \delta X_T}$$

After development and in support of (K20):

$$pT(t \rightarrow +\infty, \Delta X) = \frac{\delta X_{Max}{}^2 - \delta X_E{}^2}{2 \cdot |X_{down}| \cdot \delta X_T} = 1$$

**2/ $\Delta X \in$ [ $-\delta X_T$ ; $-\delta X_{pre}$ [**

This part of Sector 2 (Fig K2d) falls under Zone 2 and Domain 2.

So, for t infinite:

$$pT(t \to +\infty, \Delta X) = pT_{z2}(\Delta X) + pT_{d2}(t \to +\infty, \Delta X) - pT_{s2}(t \to +\infty, \Delta X)$$

With (K17b), (K18b) and (K21):

$$pT(t \to +\infty, \Delta X) = \frac{(\delta X_{Max} + \Delta X)^2}{2 \cdot \delta X_T \cdot |X_{down}|} + \frac{\delta X_{pre}}{|X_{down}| \cdot \delta X_T} \cdot \left[ \delta X_{Max} + \frac{\delta X_{pre}}{2} + \Delta X \right]$$

$$+ \frac{(|\Delta X| - \delta X_{pre})}{|X_{down}| \cdot \delta X_T} \cdot \left[ \delta X_{Max} + \frac{\delta X_{pre}}{2} + \frac{\Delta X}{2} \right] - \frac{\delta X_E{}^2}{2 \cdot |X_{down}| \cdot \delta X_T}$$

After development and using (K20):

$$pT(t \to +\infty, \Delta X) = \frac{\delta X_{Max}{}^2 - \delta X_E{}^2}{2 \cdot |X_{down}| \cdot \delta X_T} = 1$$

##### *K.4.3 Sector 3 ≡ [ $-\delta X_{Max}$ ; $-\delta X_T$ [*

Following displacement $\Delta X$, the angular range between $\theta_{down}$ and $\theta_T$ is divided into two intervals (Fig K2e)

1/ An interval I bounded by $\theta_{down}$ and ($\theta_{up}$+ $\Delta\theta$): the heads whose lever has an angle $\theta$ including these 2 terminals were in WS before the step (Fig K2e; pale green rectangle). Option 1 is respected.

**Conditions with (K3a):**

$$\left.\begin{array}{r} p = P_{SDE\_T}(t) \\ X_1 = X_{up} + \Delta X \\ X_2 = X_{down} \end{array}\right\} \Rightarrow \begin{array}{l} \delta X_L = \delta X_{Max} + \Delta X \\ \\ X_1 + X_2 = -\delta X_E + \Delta X \end{array}$$

**Calculation of the relative tension in the interval I of Sector 3 ($pT_{s3\_I}$) with application of (K15a):**

$$pT_{s3\_I}(t,\Delta X) = P_{SDE\_T}(t)\cdot\frac{(\delta X_{Max}+\Delta X)^2}{2\cdot|X_{down}|\cdot\delta X_T} \qquad \text{(K22a)}$$

2/ An interval K bounded by ($\theta_{up}$+ $\Delta\theta$) and $\theta_T$ in which heads have initiated a WS according to the 3 modes {startF}, {startS} or {startVS} (Fig K2e; pale yellow, pale mauve and pale blue rectangles superimposed). Option 2 is retained with Case 1 where the occurrence of {startF} is possible.

**Conditions with (K3b):**

$$\left.\begin{array}{r} p = P_{SDE\_1}(t) \\ X_1 = X_T \\ X_2 = X_{up} + \Delta X \end{array}\right\} \Rightarrow \begin{array}{l} \delta X_L = |\Delta X| - \delta X_T \\ \\ X_1 + X_2 = \Delta X \end{array}$$

**Calculation of the relative tension in the interval K of Sector 3 ($pT_{s3\_K}$) with application of (K15b):**

$$pT_{s3\_K}(t,\Delta X) = P_{SDE\_1}(t)\cdot\left(\frac{|\Delta X|-\delta X_T}{\delta X_T}\right)\cdot\left[1+\frac{\Delta X}{2\cdot|X_{down}|}\right] \qquad \text{(K22b)}$$

3/ The relative tension in Sector 3 is obtained by summing the two contributions on the two intervals I and K:

$$pT_{s3}(t,\Delta X) = pT_{s3\_I}(t,\Delta X) + pT_{s3\_K}(t,\Delta X)$$

So:

$$pT_{s3}(t,\Delta X) = P_{SDE\_T}(t)\cdot\frac{(\delta X_{Max}+\Delta X)^2}{2\cdot|X_{down}|\cdot\delta X_T} + P_{SDE\_1}(t)\cdot\left(\frac{|\Delta X|-\delta X_T}{\delta X_T}\right)\cdot\left[1+\frac{\Delta X}{2\cdot|X_{down}|}\right]$$

(K23)

#### Rise to T0

The expression (K23) is developed for t infinite:

$$pT_{s3}(t \to +\infty, \Delta X) = \frac{(\delta X_{Max} - |\Delta X|)^2 + (|\Delta X| - \delta X_T) \cdot (2 \cdot |X_{down}| - |\Delta X|)}{2 \cdot |X_{down}| \cdot \delta X_T}$$

Simplification implies:

$$pT_{s3}(t \to +\infty, \Delta X) = \frac{\delta X_{Max}{}^2 - 2 \cdot |X_{down}| \cdot \delta X_T}{2 \cdot |X_{down}| \cdot \delta X_T}$$

It is checked with equalities (K20) and (K21):

$$pT_{s3}(t \to +\infty, \Delta X) = pT_{s2}(t \to +\infty, \Delta X)$$

Sector 3 is in Zone 2 and belongs to Domain 2

So for t infinite:

$$pT(t \to +\infty, \Delta X) = pT_{z2}(\Delta X) + pT_{d2}(t \to +\infty, \Delta X) - pT_{s3}(t \to +\infty, \Delta X)$$

For the rise to T0 in Sector 2 with $\Delta X \in [ -\delta X_T ; -\delta X_{pre} [$, the following equality was demonstrated:

$$pT_{z2}(\Delta X) + pT_{d2}(t \to +\infty, \Delta X) - pT_{s2}(t \to +\infty, \Delta X) = 1$$

The association of the three previous formulations provides for Sector 3:

$$pT(t \to +\infty, \Delta X) = 1$$

##### *K.4.4 Sector 4 ≡ [ -($\delta X_T + \delta X_{pre}$) ; -$\delta X_{Max}$ [*

As a result of the hs shortening ($\Delta X$), none of the WS heads before the length step are still in WS after the step. All heads with a lever angle $\theta$ between $\theta_{down}$ and $\theta_T$ have initiated a WS according to the 3 modes, {startF}, {startS} or {startVS}. Option 2 is satisfied with Case 1 where the occurrence of {startF} is possible (Fig K2f; pale yellow, pale purple and pale blue rectangles superimposed).

**Conditions with (K3b):**

$$\left.\begin{array}{r} p = P_{SDE\_1}(t) \\ X_1 = X_T \\ X_2 = X_{down} \end{array}\right\} \Rightarrow \begin{array}{c} \delta X_L = \delta X_E \\ X_1 + X_2 = -\dfrac{\delta X_T}{2} - |X_{down}| \end{array}$$

**Calculation of the relative tension in mains 4 ($pT_{s4}$) with application of (K15b):**

$$pT_{s4}(t,\Delta X) = pT_{s4}(t) = P_{SDE\_1}(t) \cdot \frac{\delta X_E^{\,2}}{2 \cdot |X_{down}| \cdot \delta X_T} \qquad \text{(K24)}$$

#### <u>Rise to T0</u>

Sector 4 is outside Zone 2 and falls exclusively under Domain 3.

So for t infinite:

$$pT(t\to+\infty,\Delta X) = pT_{d3}(t\to+\infty,\Delta X) - pT_{s4}(t\to+\infty,\Delta X)$$

With (K18c) and (K24):

$$pT(t \to +\infty, \Delta X) = \frac{(\delta X_{Max} + \delta X_{pre} + \Delta X)^2 + (|\Delta X| - \delta X_{pre}) \cdot [2 \cdot \delta X_{Max} + \delta X_{pre} + \Delta X] - \delta X_E^{\,2}}{2 \cdot |X_{down}| \cdot \delta X_T}$$

After reduction and using (K20):

$$pT(t \to +\infty, \Delta X) = \frac{\delta X_{Max}^{\,2} - \delta X_E^{\,2}}{2|X_{down}| \cdot \delta X_T} = 1$$

##### *K.4.5 Sector 5 ≡ [ -($\delta X_{Max}$+ $\delta X_{pre}$ ) ; -($\delta X_T$ + $\delta X_{pre}$ ) [*

Following displacement ΔX, the angular range between $\theta_{down}$ and $\theta_T$ is divided into 2 intervals (Fig K2g).

1/ An interval I bounded by $\theta_{down}$ and ($\theta_{up}$+ $\delta\theta_{pre}$ + Δθ): the heads whose lever has an angle θ between these 2 terminals have initiated a WS according to the 3 modes, {startF}, {startS} or {startVS}, (Fig K2f ; pale yellow, pale mauve and pale blue rectangles superposed). Option 2 is agreed with Case 1 where the occurrence of {startF} is possible.

**Conditions with (K3b):**

$$\left.\begin{array}{c} p = P_{SDE\_1}(t) \\ X_1 = X_{up} + \delta X_{pre} + \Delta X \\ X_2 = X_{down} \end{array}\right\} \Rightarrow \begin{array}{c} \delta X_L = \delta X_{Max} + \delta X_{pre} + \Delta X \\ \\ X_1 + X_2 = -\delta X_E + \delta X_{pre} + \Delta X \end{array}$$

**Calculation of the relative tension in the interval I of Sector 5 ($pT_{s5\_I}$) with application of (K15b):**

$$pT_{s5\_I}(t,\Delta X) = P_{SDE\_T}(t) \cdot \frac{(\delta X_{Max} + \delta X_{pre} + \Delta X)^2}{2 \cdot |X_{down}| \cdot \delta X_T} \qquad \text{(K25a)}$$

2/ An interval K bounded by ($\theta_{up}$+ $\delta\theta_{pre}$ + Δθ) and $\theta_T$ in which heads have initiated a WS according to the only 2 modes {startS} or {startVS} (Fig K2f; pale mauve and pale blue rectangles superimposed). Option 2 is introduced with Case 2 where the occurrence of {startF} is impossible.

**Conditions with (K3c):**

$$\left.\begin{array}{c} p = P_{SDE\_2}(t) \\ X_1 = X_T \\ X_2 = X_{up} + \delta X_{pre} + \Delta X \end{array}\right\} \Rightarrow \begin{array}{c} \delta X_L = |\Delta X| - \delta X_T - \delta X_{pre} \\ \\ X_1 + X_2 = \delta X_{pre} + \Delta X \end{array}$$

**Calculation of the relative tension in the interval I of Sector 5 ($pT_{s5\_K}$) with application of (K15c):**

$$pT_{s5\_K}(t,\Delta X) = P_{SDE\_2}(t) \cdot \left( \frac{|\Delta X| - \delta X_T - \delta X_{pre}}{\delta X_T} \right) \cdot \left[ 1 + \frac{\delta X_{pre} + \Delta X}{2 \cdot |X_{down}|} \right] \qquad \text{(K25b)}$$

3/ The sum of the contributions on the 2 intervals I and K gives the relative tension in Sector 5:

$$pT_{s5}(t,\Delta X) = pT_{s5\_I}(t,\Delta X) + pT_{s5\_K}(t,\Delta X)$$

So:

$$pT_{s5}(t,\Delta X) = P_{SDE\_1}(t) \cdot \frac{(\delta X_{Max} + \delta X_{pre} + \Delta X)^2}{2 \cdot |X_{down}| \cdot \delta X_T} + P_{SDE\_2}(t) \cdot \left( \frac{|\Delta X| - \delta X_T - \delta X_{pre}}{\delta X_T} \right) \cdot \left[ 1 + \frac{\delta X_{pre} + \Delta X}{2 \cdot |X_{down}|} \right]$$

(K26)

#### Rise towards T0

Sector 5 is out of Zone 2 and belongs to Domain 3. So, for t infinite:

$$pT(t\to+\infty,\Delta X) = pT_{d3}(t\to+\infty,\Delta X) - pT_{s5}(t\to+\infty,\Delta X)$$

With (K18c) and (K26):

$$pT(t\to+\infty,\Delta X)=\frac{\left(\delta X_{Max}+\delta X_{pre}+\Delta X\right)^2}{2\cdot|X_{down}|\cdot\delta X_T}+\frac{\left(|\Delta X|-\delta X_{pre}\right)}{|X_{down}|\cdot\delta X_T}\cdot\left[\delta X_{Max}+\frac{\delta X_{pre}}{2}+\frac{\Delta X}{2}\right]$$
$$-\frac{\left(\delta X_{Max}+\delta X_{pre}+\Delta X\right)^2}{2\cdot|X_{down}|\cdot\delta X_T}-\left(\frac{|\Delta X|-\delta X_T-\delta X_{pre}}{\delta X_T}\right)\cdot\left[1+\frac{\delta X_{pre}+\Delta X}{2\cdot|X_{down}|}\right]$$

After development:

$$pT(t\to+\infty,\Delta X)=\frac{\left(|\Delta X|-\delta X_{pre}\right)}{2\cdot|X_{down}|\cdot\delta X_T}\cdot\left[2\cdot\delta X_{Max}+\delta X_{pre}+\Delta X\right]-\left(\frac{|\Delta X|-\delta X_T-\delta X_{pre}}{\delta X_T}\right)\cdot\left[\frac{2\cdot|X_{down}|+\delta X_{pre}+\Delta X}{2\cdot|X_{down}|}\right]$$

And after eliminating the ΔX-factors:

$$pT(t\to+\infty,\Delta X)=\frac{-\delta X_{pre}\cdot\delta X_T+\delta X_T\left(2\cdot|X_{down}|+\delta X_{pre}\right)}{2\cdot|X_{down}|\cdot\delta X_T}=1$$

##### *K.4.6 Sector 6 ≡ [ -150 nm ; -($\delta X_{Max}$+ $\delta X_{pre}$) [*

All heads with a lever angle θ between $\theta_{down}$ and $\theta_T$ have initiated a WS according to the 2 modes {startS} or {startVS} (Fig K2h; pale mauve and pale blue rectangles superimposed). Option 2 is posed with Case 2 where the occurrence of {startF} is impossible.

**Conditions with (K3c):**

$$\left.\begin{array}{c} p=P_{SDE\_2}(t)\\ X_1=X_T\\ X_2=X_{down}\end{array}\right\}\Rightarrow\begin{array}{c}\delta X_L=\delta X_E\\ X_1+X_2=-\frac{\delta X_T}{2}-|X_{down}|\end{array}$$

**Calculation of the tension in line 6 (pTs6) with application of (K15c):**

$$pT_{s6}(t,\Delta X)=pT_{s6}(t)=P_{SDE\_2}(t)\cdot\left(\frac{\delta X_E{}^2}{2\cdot|X_{down}|\cdot\delta X_T}\right)\qquad\text{(K27)}$$

The tension in Sector 6 is independent of step ΔX.

#### Rise to T0

Sector 4 is exclusively in Domain 4. So, for t infinite:

$$pT(t\to+\infty,\Delta X) = pT_{d4}(t\to+\infty,\Delta X) - pT_{s6}(t\to+\infty,\Delta X)$$

With (K18d) and (K27) using (K20):

$$pT(t\to+\infty,\Delta X)=\frac{\delta X_{Max}{}^2}{2\cdot|X_{down}|\cdot\delta X_T}-\frac{\delta X_E{}^2}{2\cdot|X_{down}|\cdot\delta X_T}=1$$

##### *Conclusion of the paragraph*

After an infinite time, i.e. after several dozen milliseconds under classical experimental conditions, the relative tension exerted at the 2 ends of the muscle fiber tends towards 1 and this in all cases. Consequently, the absolute tension exerted at the end of a myofibril tends towards T0, a logical conclusion because after a shortening of the fiber, the spatial distribution of the angular positions θ of the levers belonging to the WS heads converges temporally towards a uniform law on the interval $\delta\theta_T$ in any hs: this probability density characterizes the stable equilibrium of the isometric tetanus plateau (see accompanying Papers 3 and 4).

##### *Note*

For length steps less than -6 nm, the duration of phase $1(\tau_{p1})$ is experimentally increased to avoid slack phenomenon. In the example in Fig J9 illustrating paragraph J.8 of Supplement S4.J to Paper 4 where $\tau_{p1}$=1ms, it is noted that WS initiation may occur during phase 1. To account for this observation, the two time parameters $\tau_{startF}$ and $\tau_{preS}$ relating to the events {startF} and {startS} present in equations (K1a) and (K1b) are slightly modified for $\Delta X \leq -9$ nm:

- $\tau_{startF}$ will be reduced with 0.1 or 0.2 ms
- $\tau_{preS}$ will be reduced by the increased $\tau_{p1}$ value

#### K5 Master equation of the model

At the end of phase 1 (t=0), the stimulated fiber is in isometric conditions and the relative tension (pT), a function of time (t) and the length step (ΔL) corresponding to the shortening (ΔX) of any half-sarcomere of the fiber, results from the sum of 5 contributions:

$$\mathbf{pT(t,\Delta X) = pT1(\Delta X) + \Delta pT_{Relax}(t,\Delta X) + \Delta pT_{WSstart}(t,\Delta X) - \Delta pT_{FDE}(t,\Delta X) + \Delta pT_{SDE}(t,\Delta X)} \tag{K28}$$

where **pT1** is the tension at the end of phase 1 corresponding to time t=0

**$\Delta pT_{Relax}$** is the positive contribution due to the phenomenon of rapid relaxation with elimination of viscosity forces

**$\Delta pT_{WSstart}$** is the positive contribution due to the new heads that initiate a WS in the areas freed by the shortening

**$\Delta pT_{FDE}$** is the negative contribution due to the rapid detachment of heads in *down* state

**$\Delta pT_{SDE}$** is the negative contribution due to the slow detachment of the WS heads whose levers have an θ angle between $\theta_{up}$ and $\theta_T$ after the length step

The 5 terms of the right-hand member of (K28) are explained in the following sub-paragraphs.

##### *K.5.1 Relative tension at the end of phase 1 ($pT1_{Elas+Visc}$)*

The tension drop measured at the extremity of the fiber at the end of phase 1 of a length step comes from two sources:

1/ the reduction of the original elastic force following the rotation (Δθ) of the levers S1b belonging to the heads previously in WS state

2/ the braking imposed by viscosity

The sum of these two components leads to the expression (20) established in Paper 4 and reproduced below:

$$pT1 = \left[1 - P_{Elas+Visc}\right] \cdot \left(1 + \chi_{z1} \cdot K_{Z1} \cdot \overline{\Delta X}\right) + P_{Elas+Visc} \cdot \left[\chi_{z2} \cdot \left(\delta X_{Max} + K_{Z2} \cdot \overline{\Delta X}\right)\right] \tag{K29}$$

where $P_{Elas+Visc}$I is a weight varying between 0 and 1 such that:

$$P_{Elas+Visc} = \mathbf{1}_{[Bz2_{min};Bz1_{Max}[}\left(\overline{\Delta X}\right) + \left(\frac{\overline{\Delta X} - Bz1_{min}}{Bz1_{Max} - Bz1_{min}}\right) \cdot \mathbf{1}_{[Bz1_{Max};Bz1_{min}]}\left(\overline{\Delta X}\right)$$

where $\overline{\Delta X}$ is the average hs shortening which is equal by definition to:

$$\overline{\Delta X} = \frac{\sum_{h=1}^{N_{hs}} \Delta X_h}{N_{hs}} = \frac{\Delta L}{N_{hs}}$$

where $N_{hs}$ is the number of hs per myofibril; $\Delta X_h$ is the shortening of hs n° h; ΔL is the shortening of the fiber.

and where all other terms present are worded in Supplement S4.J of Paper 4.

Viscosity influences the length of the hs shortenings at the end of phase 1; see equation (J14) and Fig J4 in Supplement S4.J of Paper 4. In the tests, the lengths of the hs are generally measured within a segment located in the middle of the myofibril for which equality $\Delta X = \overline{\Delta X}$ is verified (Fig J4). Using the experimental data extracted from the frog fibers, calculations show (see the 2 cases examined in Paper 4 and Supplement S4.J) that the extreme values of hs shortening do not deviate significantly from the mean value by remaining below 10% of $\overline{\Delta X}$. So we use the formula (K29) by replacing $\overline{\Delta X}$ with ΔX.

##### *K.5.2 Contribution due to the abolition of viscosity forces ($\Delta pT_{Relax}$)*

A calculation of the tension at the end of phase 1 in the absence of viscosity ($pT1_{Elas}$) is formulated in (11) in Paper 4 and repeated below:

$$pT1_{Elas} = [1 - P_{Elas}] \cdot (1 + \chi_{z1} \cdot \Delta X) + P_{Elas} \cdot \chi_{z2} \cdot (\delta X_{Max} + \Delta X) \qquad \text{(K30)}$$

where $P_{Elas}$ is a weight equal to 0 or 1, such that:

$$P_{Elas} = \mathbf{1}_{[-\delta X_{Max};-\delta X_{z1}[}(\Delta X)$$

and where all the terms present are listed in paragraph I.4 of Supplement S4.I to Paper 4.

The end of phase 1 is the moment when the fiber is simultaneously shortened at constant speed and is already in isometry (Fig 1a of Paper 4). After phase 1, the transition between shortening at a constant speed where the viscosity forces act and isometry where the viscosity forces cancel out causes a relaxation phenomenon resulting in the rise from pT1 to $pT1_{Elas}$. It is possible to isolate the phenomenon of relaxation in the absence of any other action when the fiber is in *rigor mortaris* or at rest. Such examples are observed for various length steps in the following figures: Fig 3B from [1], Fig 1B from [2], Fig 2B from [3], Figs 1A and 1B from [4] and Fig 6 from [5]. In each of these cases, a simple exponential models the phenomenon of relaxation. An instantaneous weight ($p_{Relax}$) is introduced such that:

$$p_{Relax}(t) = e^{-\frac{t}{\tau_{Relax}}} \qquad \text{(K31)}$$

where $\tau_{Visc}$ is a time constant characteristic of relaxation in accordance with the values of the examples cited; the order of magnitude of $\tau_{Relax}$ is half a millisecond (Table 1 of Paper 5).

It is checked with (K31):

$$p_{Relax}(0) = 1$$
$$p_{Relax}(t > 4\tau_{Relax}) \approx 0$$

The positive contribution to the tension caused by the disappearance of viscosity forces ($\Delta pT_{Relax}$) is formulated from equations (K29) and (K30) associated with (K31):

$$\Delta pT_{Relax}(t, \Delta X) = (1 - p_{Relax}(t)) \cdot [\, pT1_{Elas}(\Delta X) - pT1(\Delta X)] \qquad \text{(K32)}$$

*K.5.3 Positive contribution due to the initiation of new heads towards the WS state*

The initiation of a WS corresponds to the {WSstart} event which is divided into 3 fast, slow or very slow initiation events, i.e. the 3 modes, {startF}, {startS} or {startVS}. Two instantaneous probabilities ($P_{WS1}$ and $P_{WS2}$) are presented in (K1a) and (K1b) depending on whether the occurrence of {startF} is possible or not. From these two probabilities, we define 5 weights that vary between 0 and 1 and which are a function of the numbered domain from 1 to 4 in which the tension is calculated:

$$P_{d1}(t,\Delta X) = P_{WS1}(t).\mathbb{1}_{[-\delta X_{pre};0[}(\Delta X) \tag{K33a}$$

$$P_{d2\_WS1}(t,\Delta X) = P_{WS1}(t).\mathbb{1}_{[-\delta X_{Max};-\delta X_{pre}[}(\Delta X) \tag{K33b}$$

$$P_{d2\_d3}(t,\Delta X) = P_{WS2}(t).\mathbb{1}_{[-(\delta X_{Max}+\delta X_{pre});-\delta X_{pre}[}(\Delta X) \tag{K33c}$$

$$P_{d3\_WS1}(t,\Delta X) = P_{WS1}(t).\mathbb{1}_{[-(\delta X_{Max}+\delta X_{pre});-\delta X_{Max}[}(\Delta X) \tag{K33d}$$

$$P_{d4}(t,\Delta X) = P_{WS2}(t).\mathbb{1}_{[-150nm;-(\delta X_{Max}+\delta X_{pre})[}(\Delta X) \tag{K33e}$$

The positive contribution to the instantaneous relative tension generated by the heads initiating a WS ($\Delta pT_{WSstart}$) is based on equations (K7) to (K11) established in paragraph K.2 and in support of the weights determined in (K33a) to (K33e):

$$\begin{aligned}
\Delta\, pT_{WSstart}(t,\Delta X) = {} & P_{d1}(t,\Delta X)\cdot\frac{|\Delta X|\cdot\left[\delta X_{Max}+\frac{\Delta X}{2}\right]}{|X_{down}|\cdot\delta X_T} \\
& + P_{d2_WS1}(t,\Delta X)\cdot\frac{\delta X_{pre}}{|X_{down}|\cdot\delta X_T}\cdot\left[\delta X_{Max}+\frac{\delta X_{pre}}{2}+\Delta X\right] \\
& + P_{d2_d3}(t,\Delta X)\cdot\frac{\left(|\Delta X|-\delta X_{pre}\right)}{|X_{down}|\cdot\delta X_T}\cdot\left[\delta X_{Max}+\frac{\delta X_{pre}}{2}+\frac{\Delta X}{2}\right] \\
& + P_{d3_WS1}(t,\Delta X)\cdot\frac{\left(\delta X_{Max}+\delta X_{pre}+\Delta X\right)^2}{2\cdot|X_{down}|\cdot\delta X_T} \\
& + P_{d4}(t,\Delta X)\cdot\frac{\left(\delta X_{Max}\right)^2}{2\cdot|X_{down}|\cdot\delta X_T}
\end{aligned} \tag{K34}$$

*K.5.4 Negative contribution due to the fast detachment event {FastDE}*

This negative contribution is developed in paragraph K.3 with equation (K12) as following:

$$\Delta\, pT_{FDE}(t,\Delta X) = P_{FDE}(t)\cdot\left[\frac{(\delta X_{Max}+\Delta X)^2}{2\cdot\delta X_T\cdot|X_{down}|} - \chi_{z2}\cdot(\delta X_{Max}+\Delta X)\right] \qquad \text{(K35)}$$

where $P_{FDE}$ is a weight ranging from 0 to 1, such that:

$$P_{FDE}(t) = \left(1 - e^{-\frac{t-\tau_{preFDE}}{\tau_{FDE}}}\right).\mathbb{1}_{[-\delta X_{Max};-\delta X_{z1}]}(\Delta X)$$

*K.5.5 Negative contribution due to the event slow detachment {SlowDE}*

The slow detachment of a WS myosin head is presented in paragraph K.4. Three instantaneous probabilities ($P_{SDE\_T}$ , $P_{SDE\_1}$ , $P_{SDE\_2}$) are given in (K3a), (K3b) and (K3c), respectively. With these 3 probabilities, we define 6 weights which are function of the indexed sector from 1 to 6 where the tension is calculated:

$$P_{s1}(t,\Delta X) = P_{SDE\_T}(t).\mathbb{1}_{[-\delta X_E;0[}(\Delta X) \qquad \text{(K36a)}$$

$$P_{s3\_T}(t,\Delta X) = P_{SDE\_T}(t).\mathbb{1}_{[-\delta X_{Max};-\delta X_T[}(\Delta X) \qquad \text{(K36b)}$$

$$P_{s3\_DE1}(t,\Delta X) = P_{SDE\_1}(t).\mathbb{1}_{[-\delta X_{Max};-\delta X_T[}(\Delta X) \qquad \text{(K36c)}$$

$$P_{s5\_DE1}(t,\Delta X) = P_{SDE\_1}(t).\mathbb{1}_{[-(\delta X_{Max}+\delta X_{pre});-(\delta X_T+\delta X_{pre})[}(\Delta X) \qquad \text{(K36d)}$$

$$P_{s5\_DE2}(t,\Delta X) = P_{SDE\_2}(t).\mathbb{1}_{[-(\delta X_{Max}+\delta X_{pre});-(\delta X_T+\delta X_{pre})[}(\Delta X) \qquad \text{(K36e)}$$

$$P_{s2,s4,s6}(t,\Delta X) = P_{SDE\_T}(t).\mathbb{1}_{[-\delta X_T;-\delta X_E[}(\Delta X) + P_{SDE\_1}(t).\mathbb{1}_{[-(\delta X_T+\delta X_{pre});-\delta X_{Max}]}(\Delta X) + P_{SDE\_2}(t)..\mathbb{1}_{[-150nm;-(\delta X_{Max}+\delta X_{pre})]}(\Delta X) \qquad \text{(K36f)}$$

The negative contribution to the instantaneous relative stress generated by the slowly detaching heads ($\Delta pT_{SDE}$) is based on equations (K19) to (K27) and in support of equations (K36a) to (K36f):

$$\Delta pT_{SDE}(t,\Delta X) = P_{s1}(t,\Delta X)\cdot\frac{|\Delta X|\cdot\left[\delta X_E+\frac{\Delta X}{2}\right]}{|X_{down}|\cdot\delta X_T}$$

$$+P_{s3\_T}(t,\Delta X)\cdot\frac{(\delta X_{Max}+\Delta X)^2}{2\cdot|X_{down}|\cdot\delta X_T}$$

$$+P_{s3\_DE1}(t,\Delta X)\cdot\left(\frac{|\Delta X|-\delta X_T}{\delta X_T}\right)\cdot\left[1+\frac{\Delta X}{2\cdot|X_{down}|}\right]$$

$$+P_{s5\_DE1}(t,\Delta X)\cdot\frac{(\delta X_{Max}+\delta X_{pre}+\Delta X)^2}{2\cdot|X_{down}|\cdot\delta X_T}$$

$$+P_{s5\_DE2}(t,\Delta X)\cdot\left(\frac{|\Delta X|-\delta X_T-\delta X_{pre}}{\delta X_T}\right)\cdot\left[1+\frac{\delta X_{pre}+\Delta X}{2\cdot|X_{down}|}\right]$$

$$+P_{s2,s4,s6}(t,\Delta X)\cdot\frac{\delta X_E{}^2}{2\cdot|X_{down}|\cdot\delta X_T}$$

(K37)
