## Supplementary material for "Mechanical model of muscle contraction. 5. Tension rise after phase 1 of a length step": Computer Programs relative to the calculations and plots in Paper 5.

### CP5 Computer Programs relative to the calculations and plots in Paper 5

#### CP5.1 Calculation and curve of T as a function of ΔX and t

The curve on the computer screen is done by calling the "SUB AA_REMontees_trace()" routine.

Once the data have been recovered (See values displayed in the Tables of Paper 5 and in the Excel sheets of Supplement DA5, the experimental tension from published articles and the theoetrical tension curves from model equations are plotted as a function of the time and the hs shortening (Fig CP5.1).

A regression is performed between experimental and theoretical tensions.

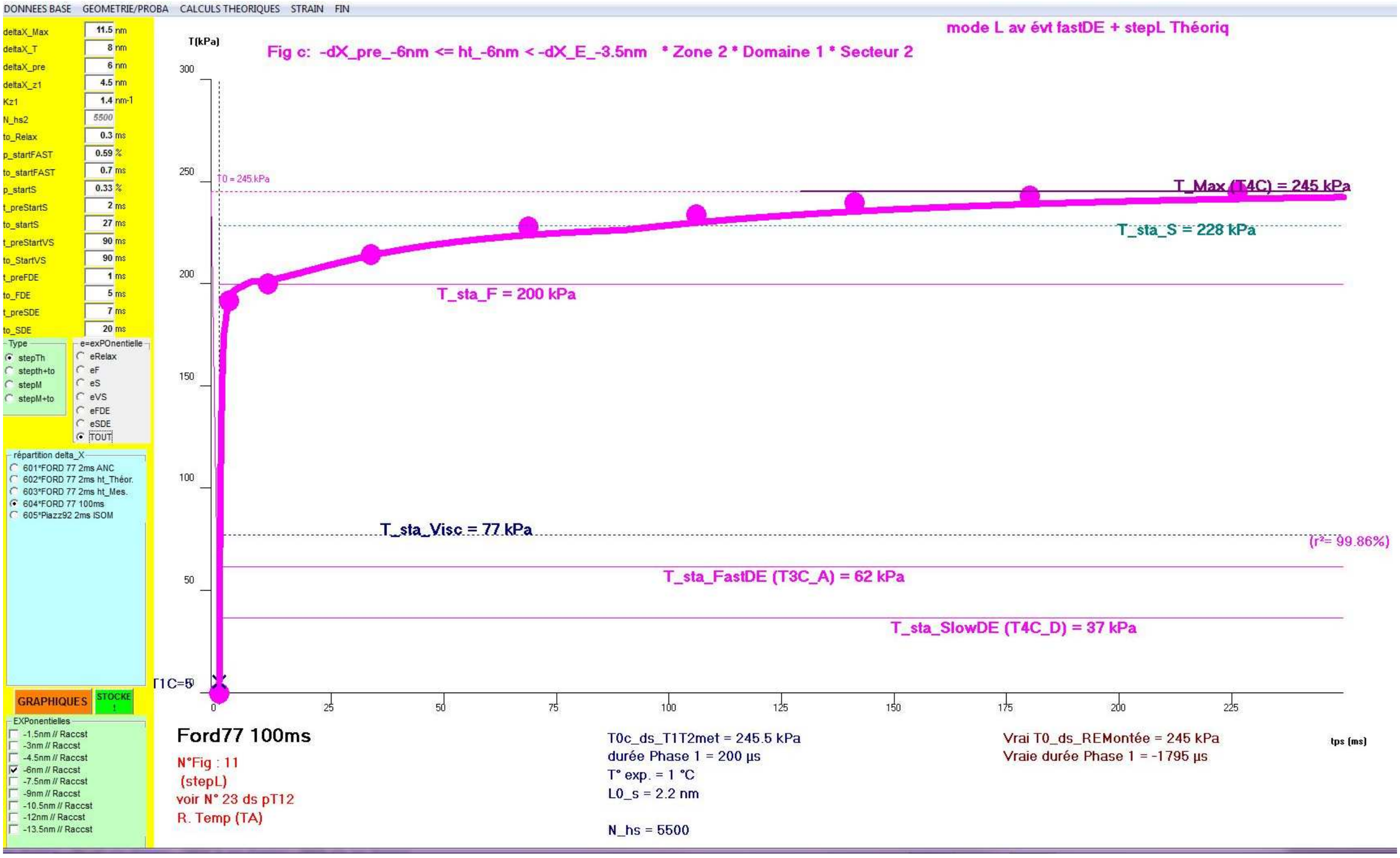


**Fig CP5.1 Screenshot after starting the the "Sub AAPH_REMontee_trace()" routine.**

#### Sub AA_REMontees_trace(iX As Integer)

```
e_pt = 25 '15 si toutes les step sur la m courbe
e_tt = 8 '6 si tout sur m courbe
ToupT = 1 'par defaut
zfm = iX + FraX.Tag
T1T2met.Seek "=", zfm
ToupT = 1 'T en kPa
'ToupT = 2 'pT en %
Oesc(4).Enabled = IIf(Frtyp.Tag < 3, -1, 0) 'pas de FastDE en mode Amorti
With CD_GL: .Top = Height * 0.65 : .Left = Width * 0.91 : .Visible = -1 : End With
With CD_BA : .Top = Height * 0.75 : .Left = Width * 0.91 : .Visible = -1 : End With
With Frpres : .Top = Height * 0.35: .Left = Width * 0.9 : .Visible = -1 : End With
For t = 0 To 4
      With Opres(t)
         If t < 4 Then
           .Visible = -1
           .Caption = Choose(t + 1, "TOUT", "PTS SANS légende", "PAS de PTS AVEC légende", "PAS de PTS SANS légende", "", "")
         Else
            .Visible = 0
         End If
      End With
   Next
End If
```

##### AA_REMontees_trace_prepa zfm

```
Npts = 100
coul_2 = QBColor(1)

For t = 1 To FrpTouT.Tag
If ChT1T2(t) = 1 Then
       REMo.Seek "=", zfm, t
       N_hs = T1T2met("n_hs")
       coul = QBColor(REMo("coul_pt"))
   '** depart ISOMETREI TETANIQUE TOc
       T0c = REMo("T0")
       If ToupT = 1 Then
         yav = (T0c - yn) * 100 / (yx - yn)
       Else
        yav = (1 - yn) * 100 / (yx - yn)
       End If
        xav = (REMo("tps_T0") - xn) * 100 / (xx - xn)
        O.DrawWidth = 2
        O.Line (0, yav)-(xav, yav), coul
        If N_STEP = 1 Then
            O.DrawStyle = 2
            O.DrawWidth = 1
            O.Line (xav, yav)-(100, yav), coul
            O.DrawStyle = 0
            If Frpres.Tag = 0 Or Frpres.Tag = 2 Then 'légendes
               O.CurrentX = 0.5: O.CurrentY = yav + IIf(REMo("stepL") > 0, -1, 3)
               O.ForeColor = coul
               O.Print "T0 = " + Format(T0c, "###.#") + "kPa"
          End If
  '** PHASE 1: 1 droite jusqu'à (t=0,T=T1M)
         O.DrawWidth = 1
         If REMo("tps_slack") < 0 Then
            'd'abord segment droite de T0 à slack
            xap = (REMo("tps_slack") - xn) * 100 / (xx - xn)
            yap = 0
            O.Line (xav, yav)-(xap, yap), coul
             xav = xap: yav = 0
         End If
         xap = -xn * 100 / (xx - xn)
         yap = REMo("T1M") * 100 / (yx - yn)
          O.Line (xav, yav)-(xap, yap), coul
   'les points
        If (ressort.Tag = "REMontee_stepL_OK" And Frpres.Tag < 2) Or Left(ressort.Tag, 16) = "REMontee_stepL_p" Then
           O.DrawWidth = e_pt
           For i = 0 To REMo("N_pts")
              xav = (REMo("tps_" + Format(i)) - xn) * 100 / (xx - xn)
              ymi = REMo("Tin_" + Format(i))
              yav = (ymi - yn) * 100 / (yx - yn)
```

```
        O.PSet (xav, yav), coul
      Next
    End If
  'trait +nom du stepL si pas de légende
    If ressort.Tag = "REMontee_stepL_OK" And (Frpres.Tag = 1 Or Frpres.Tag = 3) Then
      O.DrawWidth = 5
      xav = 85
      yav = 40 - t * 4
      O.Line (xav, yav)-(xav + 4, yav), coul
      O.CurrentX = xav + 5
      O.CurrentY = yav + 2
      O.ForeColor = coul
       O.FontBold = -1: O.FontSize = 14
      tx1 = IIf(REMo("stepL") > 0, "+", "") + Format(REMo("stepL")) + " nm"
      O.Print tx1
    End If
  End If
'***etap 2 on trace le/les expon
  AA_REMontees_trace_D_stepl_OK1 'prepa des données
  AA_REMontees_trace_D_stepl_OK2 'tracé des exponentielles
  If Fresc.Tag = 6 Then 'regression des tensions
      REMr2.Index = "temps"
      sX = 0: sY = 0: sXX = 0: sYY = 0: sXY = 0
      For i = 0 To REMo("N_pts")
        xav = REMo("tps_" + Format(i))
       'mesuré
        yav = REMo("Tin_" + Format(i))
       'theorique
        REMr2.Seek ">=", xav: If REMr2.NoMatch Then Stop
        If xav = REMr2("temps") Then
          yap = REMr2("pTt")
          REMr2.Edit: REMr2("pTm") = yav: REMr2.Update
        ElseIf xav < REMr2("temps") Then 'faut interpoler
          xmi = REMr2("temps")
          ymi = REMr2("pTt")
          REMr2.MovePrevious
          yap = REMr2("pTt") + (ymi - REMr2("pTt")) * (xav - REMr2("temps")) / (xmi - REMr2("temps"))
          'REMr2.AddNew: REMr2("pTm") = yav: REMr2.Update
          REMr2.AddNew
           REMr2("cas") = zfm
           REMr2("step") = t
           REMr2("pt") = REMr2.RecordCount + 1
           REMr2("temps") = xav
           REMr2("pTm") = yav
           REMr2("pTt") = yap
         REMr2.Update
        Else
          Stop
        End If
       'calcul
         sX = sX + yav: sXX = sXX + yav ^ 2
         sY = sY + yap: sYY = sYY + yap ^ 2
         sXY = sXY + yav * yap
      Next 'i
      sN = REMo("N_pts") + 1
      p_regL = sXY / sXX
      r_regL = 1 - (sYY - 2 * p_regL * sXY + sXX * p_regL ^ 2) / (sYY - sY * sY / sN)
      REMo.Edit
       REMo("r2") = r_regL
       REMo("pente") = p_regL
      REMo.Update
      'on ecrit
       O.ForeColor = coul: O.FontBold = -1
      O.CurrentX = 97
      O.CurrentY = 42 - t * 4
      tx1 = "(r²= " + Format(r_regL * 100, "###.##") + "%)"
     O.Print tx1
    End If
End If ' case cochée  ChT1T2(t) = 1 Then
Next ' t =step
End Sub
```

#### Sub AA_REMontees_trace_prepa(Ino As Integer)

```
impscale X0, Y0 + 130, X0 + larg, Y0 - 10 '14
    vx = "tps (ms)"    'le xx est calculé d'apres les valeurs ds Sub_AA_REMontees_T1T2_step
  Select Case xx
     Case Is <= 2: xt = 0.1
     Case Is <= 20: xt = 1
     Case Is <= 60: xt = 5
     Case Is <= 120: xt = 10
      Case Is <= 500: xt = 25
     Case Is <= 1000: xt = 100
     Case Else: xt = 500
    End Select
        If ToupT = 2 Then
        vy = "pT(%)": yx = 1.2: yt = 0.2
    Else
        vy = "T(kPa)": yt = 50
    End If
    impenvi
    xav = -xn * 100 / (xx - xn)

    'ligne ne pointille du temps_zero(apres remontée T2visc)
    O.ForeColor = QBColor(0)
    O.DrawStyle = 2
    O.Line (xav, 0)-(xav, 100) ', QBColor(12)
    O.DrawStyle = 0

End Sub
```

#### Sub AA_REMontees_trace_D_stepl_OK1()

```
If IsNull(REMo("T_deb")) Then AA_REMontees_trace_D_stepl_OK_rempli 'on ne le fait que la 1ere fois
If N_STEP = 1 Then
    L_WS = Val(TAM(0).Text)
    aX = Val(TAM(1).Text)
    L_preWS = Val(TAM(2).Text)
    dXz1 = Val(TAM(3).Text)
    KqnZ1 = Val(TAM(4).Text) ' =nu_visc
    to_Relax = Val(TAM(6).Text)
    P_startF = Val(TAM(7).Text)
    to_startF = Val(TAM(8).Text)
    P_startS = Val(TAM(9).Text)
    t_preS = Val(TAM(10).Text)
    to1_startS = Val(TAM(11).Text)
    t_preVS = Val(TAM(12).Text)  '70-150 ms
    to1_startVS = Val(TAM(13).Text) '100 ms
    t_pre_FastDE = Val(TAM(14).Text) 't_deb_phase2=5 ms ?
    to_FastDE = Val(TAM(15).Text)    '3 à 6 ms ?
      t_pre_SlowDE = Val(TAM(16).Text)  't>10 ms ?
    to_SlowDE = Val(TAM(17).Text)     '= to_startS = 30 ms ?

  Else 'on recupere dans la table directement
     L_WS = REMo("L_WS")
     aX = REMo("aX")
     L_preWS = REMo("L_preWS")
     dXz1 = REMo("dXz1")
     KqnZ1 = REMo("Kz1")
     to_Relax = REMo("to_Relax")
     P_startF = REMo("P_startF")
     to_startF = REMo("to_startF")
     P_startS = REMo("P_startS")
     t_preS = REMo("t_preS")
     to1_startS = REMo("to1_startS")
     t_preVS = REMo("t_preVS")
     to1_startVS = REMo("to1_startVS")
     t_pre_FastDE = REMo("t_pre_FastDE")
     to_FastDE = REMo("to_FastDE")
    t_pre_SlowDE = REMo("t_pre_slowDE")
    to_SlowDE = REMo("to_slowDE")
  End If
TAM(5).Text = Format(N_hs)
Set SL = REMo
 tx1 = "stepL" + IIf(Frtyp.Tag < 3, "", "_Mdeb")
 ht_step = SL(tx1)
 AA_REMontees_trace_D_stepL_OK1_prepa
 'etap4: TRACES des DROITES parallele auxabscisses pour svoir sur quoi on travaille
 O.CurrentX = 65: O.CurrentY = 110
 O.FontBold = -1: O.FontSize = 14
 O.Print Choose(Frtyp.Tag, "mode L av évt fastDE + stepL Théoriq", "mode L av évt fastDE + stepL Mesuré", "Mode_A ss évt FastDE + stepL Théoriq", "Mode_A ss évt FastDE + stepL Mesuré")
If Frpres.Tag = 0 Or Frpres.Tag = 2 Then 'légendes
   tx1 = IIf(REMo("stepL") < 0, "+", "")
   'droites commentaires
    O.ForeColor = coul_2 'QBColor(1)
    O.DrawWidth = 1
    xav = -xn * 100 / (xx - xn)
  'droite T2C_vi0 (dispartion viscosité)     'T_STA_Vi = REMo("T2C_vi0")
    yav = T_STA_Vi * 100 / (yx - yn)
    O.DrawStyle = 2
    O.Line (xav, yav)-(100, yav), coul_2
    O.DrawStyle = 0
    O.CurrentX = 15: O.CurrentY = yav + 2.5: O.Print "T_sta_Visc = " + Format(Int(T_STA_Vi + 0.5)) + " kPa"
  'droite T2C_mR (Ancien modele Rapide)
    yav = T_STA_F * 100 / (yx - yn)
    O.Line (xav, yav)-(100, yav), coul
    O.ForeColor = coul
    O.CurrentX = 20: O.CurrentY = yav: O.Print "T_sta_F = " + Format(Int(T_STA_F + 0.5)) + " kPa"
   'droite T3C_A (mode Amorti)
      yav = T_sta_FastDE * 100 / (yx - yn)
      O.Line (xav, yav)-(100, yav), coul
      O.CurrentX = 40: O.CurrentY = yav: O.Print "T_sta_FastDE (T3C_A) = " + Format(Int(T_sta_FastDE + 0.5)) + " kPa"
```

```
 'droite T4C_D (mode Détaché)    'T_sta_SlowDE = REMo("T4C_D")
        yav = T_sta_SlowDE * 100 / (yx - yn)
        O.Line (xav, yav)-(100, yav), coul
        O.CurrentX = 60: O.CurrentY = yav: O.Print "T_sta_SlowDE (T4C_D) = " + Format(Int(T_sta_SlowDE + 0.5)) + " kPa"
     O.DrawWidth = 1: O.DrawStyle = 2
     yav = T_STA_S * 100 / (yx - yn)
        O.Line (xav, yav)-(100, yav), QBColor(3)
        O.ForeColor = QBColor(3)
        O.CurrentX = 80: O.CurrentY = yav + 1: O.Print "T_sta_S = " + Format(Int(T_STA_S + 0.5)) + " kPa"
      O.DrawStyle = 0
     ' droite T_STA_S=VT0c       'T_STA_VS = T_Max= REMo("T4C")  '= REMo("T4C_D") + REMo("remT2_mR") + REMo("remT4_S")
       O.DrawWidth = 2
      yav = T_MAX * 100 / (yx - yn)
        O.Line (52, yav)-(100, yav), QBColor(5)
        O.ForeColor = QBColor(5)
        O.CurrentX = 85: O.CurrentY = yav + 2.5: O.Print "T_Max (T4C) = " + Format(Int(T_MAX + 0.5)) + " kPa"
        'zonz+domaine+secteur
        If ht_step >= Xmin Then 'Fig M3b
          tx1 = "Fig b:  -dX_E_" + Format(Xmin) + "nm <= ht_" + Format(ht_step) + "nm < 0"
          xav = 1: xmi = 1: xap = 1
        ElseIf ht_step >= -L_preWS Then 'Fig M3c
          tx1 = "Fig c:  -dX_pre_" + Format(-L_preWS) + "nm <= ht_" + Format(ht_step) + "nm < -dX_E_" + Format(Xmin) + "nm"
          xav = 2: xmi = 1: xap = 2
        ElseIf ht_step >= -aX Then 'Fig M3d
          tx1 = "Fig d:  -dX_T_" + Format(-aX) + "nm <= ht_" + Format(ht_step) + "nm < -dX_pre_" + Format(-L_preWS) + "nm"
          xav = 2: xmi = 2: xap = 2
        ElseIf ht_step >= -L_WS Then 'Fig M3e
          tx1 = "Fig e:  -dX_Max_" + Format(-L_WS) + "nm <= ht_" + Format(ht_step) + "nm < -dX_T_" + Format(-aX) + "nm"
          xav = 2: xmi = 2: xap = 3
        ElseIf ht_step >= -(aX + L_preWS) Then 'Fig M3f
          tx1 = "Fig f:  -(dX_T+dX_pre)_" + Format(-aX - L_preWS) + "nm <= ht_" + Format(ht_step) + "nm < -dX_E_" + Format(-L_WS) + "nm"
          xav = 3: xmi = 3: xap = 4
        ElseIf ht_step >= -(L_WS + L_preWS) Then 'Fig M3g
          tx1 = "Fig g:  -(dX_Max+dX_pre)_" + Format(-L_WS - L_preWS) + "nm <= ht_" + Format(ht_step) + "nm < -(dX_T+dX_pre)_" + Format(-aX - L_preWS) + "nm"
          xav = 3: xmi = 3: xap = 5
        Else 'Fig M3h
          tx1 = "Fig h:  ht_" + Format(ht_step) + "nm > -(dX_Max+dX_pre)_" + Format(-L_WS - L_preWS) + "nm"
          xav = 3: xmi = 4: xap = 6
        End If
        O.ForeColor = coul 'QBColor(5)
        O.CurrentX = 5: O.CurrentY = 106
        O.Print tx1 + "    * Zone " + Format(xav) + " * Domaine " + Format(xmi) + " * Secteur " + Format(xap)
End If
   O.FontSize = 18
   O.CurrentX = -3:  O.CurrentY = -5
   O.ForeColor = QBColor(0)
   O.Print T1T2met("fm")   '+ "_" + fm("method")
   O.FontBold = -1:  O.FontSize = 12

   For i = 1 To 14
        tx1 = ""
        Select Case i
             Case 1: tx1 = "N°Fig : " + T1T2met("no_Fig")
             Case 2
               If IsNull(T1T2met("facteur")) = 0 Then
                  tx1 = tx1 + " (" + T1T2met("facteur") + ")"
                End If
             Case 3: If IsNull(T1T2met("comm")) = 0 Then tx1 = T1T2met("comm")
             Case 4: tx1 = T1T2met("l_Espece") + " (" + T1T2met("l_Musc") + ")"
             Case 5:  If IsNull(T1T2met("T0c")) = 0 Then tx1 = "T0c_ds_T1T2met = " + Format(T1T2met("T0c")) + " kPa"
             Case 6: If IsNull(T1T2met("du_p1")) = 0 Then tx1 = "durée Phase 1 = " + T1T2met("du_p1") + " µs"
             Case 7: If IsNull(T1T2met("Texp")) = 0 Then tx1 = "T° exp. = " + T1T2met("Texp") + " °C"
             Case 8: If IsNull(T1T2met("L0_s")) = 0 Then tx1 = "L0_s = " + T1T2met("L0_s") + " nm"
             Case 9:
             Case 10: tx1 = "N_hs = " + Format(N_hs)
             Case 11: If IsNull(REMo("T0")) = 0 Then tx1 = "Vrai T0_ds_REMontée = " + Format(Int(REMo("T0") * 10 + 0.5) / 10) + " kPa"
             Case 12: If IsNull(REMo("tps_T0")) = 0 Then tx1 = "Vraie durée Phase 1 = " + Format(Int(REMo("tps_T0") * 1000 + 0.5)) + " µs"
             Case 13: If IsNull(REMo("tps_slack")) = 0 Then tx1 = "durée slack = " + Format(Int(REMo("tps_slack") * 1000 + 0.5)) + " µs"
             Case 14: 'If IsNull(T1T2met("L_WS")) = 0 Then tx1 = "dX_Max = " + Format(T1T2met("L_WS"))
           End Select
```

```
      If Len(tx1) > 0 Then
        If i < 5 Then
          O.ForeColor = QBColor(12)
          O.CurrentX = -3
          O.CurrentY = -7 - i * 3
        ElseIf i < 11 Then
          O.ForeColor = QBColor(1)
          O.CurrentX = 35
          O.CurrentY = -(i - 3) * 3
        Else
          O.ForeColor = QBColor(4)
          O.CurrentX = 70
          O.CurrentY = -(i - 9) * 3
        End If
        O.Print tx1
      End If
  Next 'i
End Sub
```

#### Sub AA_REMontees_trace_D_stepL_OK1_prepa()

```
Dim remT_WSstart As Single, dimT_SDE As Single
'Rappels: ax=dX_start // L_WS=dX_Max // L_preWS=dX_pre // Xmin= -dX_end
  X1 = aX / 2      'X_up
  X2 = -X1        'X_T
  X3 = X1 - L_WS   'X_down
  Xmin = aX - L_WS '-dX_E <0
  chi_p1 = 1 / Abs(X3)
  If dXz1 < Abs(Xmin) Then
     dXz1 = Abs(Xmin)
     TAM(3).Text = Format(dXz1)
  End If
  X_Z1 = -dXz1
  chi_p2 = (1 + X_Z1 * chi_p1) / (L_WS + X_Z1)

'-------qZ1------------KqnZ1 = N_hs ^ (1 - qZ1 / 2) * coth[N_hs ^ (1 - qZ1 / 2)]
calcul_qoK (KqnZ1) '1 / 8.01) * Log(2800000 / Kqn) '1.9
qZ1 = qOK
xmi = N_hs ^ (1 - qZ1 / 2)
cH_n = (Exp(xmi) + Exp(-xmi)) / 2
sH_n = (Exp(xmi) - Exp(-xmi)) / 2
z1_min = X_Z1 / KqnZ1
z1_max = z1_min * cH_n
'-------rZ2------------
rZ2 = qZ1 - Log(chi_p1 / chi_p2) / Log(N_hs)
xmi = N_hs ^ (1 - rZ2 / 2)
cH_n = (Exp(xmi) + Exp(-xmi)) / 2
sH_n = (Exp(xmi) - Exp(-xmi)) / 2
KrnZ2 = calcul_Kqn(rZ2)
z2_min = -L_WS / KrnZ2
z2_max = z2_min * cH_n
X_T0 = z2_min
 T0c = SL("T0")
SL.Edit
'** PHASE 1
            'T1C = T1_avV = pPT1_Elas+Visc   ...... calculé'attention c'est 1 cste il faut donc récupérer T0(c)
             If ht_step > 4 Then 'stretching trop important
                        Stop ' a faire elasticit parabolique du aux S2 ?
             ElseIf ht_step >= z1_min Then 'tous les hs en ZONE 1 avec stretching compris jusqu'à 4nm ?
                        SL("T1C") = T0c * (1 + ht_step * chi_p1 * KqnZ1) '
             ElseIf ht_step >= z1_max Then 'ZONE MIXTE  hs ds zone 1 et 2
                         yav = T0c * (1 + ht_step * chi_p1 * KqnZ1)
                         yap = T0c * (Abs(X_T0) + ht_step) * chi_p2 * KrnZ2
                         poids = (ht_step - z1_min) / (z1_max - z1_min)
                         SL("T1C") = (1 - poids) * yav + poids * yap
              ElseIf ht_step >= X_T0 Then
                          SL("T1C") = T0c * (Abs(X_T0) + ht_step) * chi_p2 * KrnZ2
              Else
                        SL("T1C") = 0
              End If
             If SL("T1C") < 0 Then SL("T1C") = 0

 '** PHASE 2/3/4
        If ht_step >= X_Z1 Then
                        SL("T2C_vi0") = T0c * (1 + ht_step * chi_p1)
        ElseIf ht_step >= -L_WS Then
                        SL("T2C_vi0") = T0c * (L_WS + ht_step) * chi_p2
        Else
                        SL("T2C_vi0") = 0
        End If

        If ht_step > 0 Then 'steching
                    SL("remT_F") = 0
                    SL("remT_S") = 0
                    SL("remT_VS") = 0
         ElseIf ht_step >= -L_preWS Then 'domaine 1
                      sX = T0c * (chi_p1 / aX) * Abs(ht_step) * (L_WS + ht_step / 2)
                     SL("remT_F") = P_startF * sX
                     SL("remT_S") = P_startS * sX
                     SL("remT_VS") = (1 - P_startH) * sX
```

```
  ElseIf ht_step >= -L_WS Then 'domaine 2
           sX = T0c * (chi_p1 / aX) * L_preWS * (L_WS + L_preWS / 2 + ht_step)
           sY = T0c * (chi_p1 / aX) * (Abs(ht_step) - L_preWS) * (L_WS + L_preWS / 2 + ht_step / 2)
           SL("remT_F") = P_startF * sX
           SL("remT_S") = P_startS * sX + P_startH * sY
           SL("remT_VS") = (1 - P_startH) * (sX + sY)
  ElseIf ht_step >= -(L_WS + L_preWS) Then 'domaine 3
           sX = T0c * (0.5 * chi_p1 / aX) * (L_WS + L_preWS + ht_step) ^ 2
           sY = T0c * (chi_p1 / aX) * (Abs(ht_step) - L_preWS) * (L_WS + L_preWS / 2 + ht_step / 2)
           SL("remT_F") = P_startF * sX
           SL("remT_S") = P_startS * sX + P_startH * sY
           SL("remT_VS") = (1 - P_startH) * (sX + sY)
   Else 'Domaine 4 independante de DX
           sX = T0c * chi_p1 * L_WS ^ 2 / (2 * aX)
           SL("remT_F") = 0
           SL("remT_S") = P_startH * sX
           SL("remT_VS") = (1 - P_startH) * sX
    End If

 'T3C_A ...calculé pour Evt_FastDE avec mode Amorti (t_p1> 2ms)
    If ht_step > 0 Then 'steching
          SL("T3C_A") = 0  2
    ElseIf ht_step >= X_Z1 Then 'ou Xmin ?
           SL("T3C_A") = T0c * (1 + ht_step * chi_p1)
     ElseIf ht_step >= -L_WS Then
          SL("T3C_A") = T0c * (0.5 * chi_p1 / aX) * (L_WS + ht_step) ^ 2
     Else
           SL("T3C_A") = 0
     End If

'T4C_D ...calculé pour Evt_SlowDE avec mode Détaché Lent
    If ht_step > 0 Then 'steching
           SL("dimT_Te_SDE") = 0 'Tetanos puis SDE
           SL("dimT_F_SDE") = 0 'F=startF puis SDE
           SL("dimT_S_SDE") = 0 'S=startS puis SDE
           SL("dimT_VS_SDE") = 0 'VS=startVS puis SDE
     ElseIf ht_step >= Xmin Then 'Secteur 1
           SL("dimT_Te_SDE") = T0c * (chi_p1 / aX) * Abs(ht_step) * (Abs(Xmin) + ht_step / 2)
           SL("dimT_F_SDE") = 0
           SL("dimT_S_SDE") = 0
           SL("dimT_VS_SDE") = 0
    ElseIf ht_step >= -aX Then 'Secteur 2
             SL("dimT_Te_SDE") = T0c * (0.5 * chi_p1 / aX) * Xmin ^ 2
           SL("dimT_F_SDE") = 0
           SL("dimT_S_SDE") = 0
           SL("dimT_VS_SDE") = 0
   ElseIf ht_step >= -L_WS Then 'Secteur 3
           SL("dimT_Te_SDE") = T0c * (0.5 * chi_p1 / aX) * (L_WS + ht_step) ^ 2
             sX = T0c * (Abs(ht_step) / aX - 1) * (1 + chi_p1 * ht_step / 2)
           SL("dimT_F_SDE") = P_startF * sX
           SL("dimT_S_SDE") = P_startS * sX
           SL("dimT_VS_SDE") = (1 - P_startH) * sX
    ElseIf ht_step >= -(aX + L_preWS) Then 'Secteur 4
          SL("dimT_Te_SDE") = 0
           sX = T0c * (0.5 * chi_p1 / aX) * Xmin ^ 2
           SL("dimT_F_SDE") = P_startF * sX
           SL("dimT_S_SDE") = P_startS * sX
           SL("dimT_VS_SDE") = (1 - P_startH) * sX
    ElseIf ht_step >= -(L_WS + L_preWS) Then 'Secteur 3
           SL("dimT_Te_SDE") = 0
           sX = T0c * (0.5 * chi_p1 / aX) * (L_WS + L_preWS + ht_step) ^ 2
           sY = T0c * (Abs(ht_step) - aX - L_preWS) * (1 + chi_p1 * (L_preWS + ht_step) / 2) / aX
           SL("dimT_F_SDE") = P_startF * sX
           SL("dimT_S_SDE") = P_startS * sX + P_startH * sY
           SL("dimT_VS_SDE") = (1 - P_startH) * (sX + sY)
    Else
           SL("dimT_Te_SDE") = 0
          SL("dimT_F_SDE") = 0
          sY = T0c * (0.5 * chi_p1 / aX) * Xmin ^ 2
          SL("dimT_S_SDE") = P_startH * sY
          SL("dimT_VS_SDE") = (1 - P_startH) * sY
```

```
            End If

'etap3: bilan
    SL("T_deb") = SL("T1C")
    T_deb = SL("T_deb") 'T1_Elas+Visc

  '** Phase 2 remontee rapide  viscosité
    SL("T_STA_Vi") = SL("T2C_vi0")
    T_STA_Vi = SL("T_STA_Vi") 'apres relax T1_Elas

  '** Phase 2 startF
    If ht_step > 0 Then 'Allgt
       SL("T2C_mR") = T0c + 5 'par défaut
    Else
       SL("T2C_mR") = T_STA_Vi + SL("remT_F")
    End If
    SL("T_STA_F") = SL("T2C_mR")
    T_STA_F = SL("T2C_mR") ' Fin remontéee apres intitaioation rapide

  'Phase 3 FastDE retour vers mode Amorti
    SL("T_STA_FastDE") = SL("T3C_A")
    T_sta_FastDE = SL("T3C_A") 'T1_Elas _Parabole pares detachement

  'Phase 4 SlowDE  tete en tre TeTa_down et TeTa_T
    remT_WSstart = SL("remT_F") + SL("remT_S") + SL("remT_VS")
    dimT_SDE = SL("dimT_Te_SDE") + SL("dimT_F_SDE") + SL("dimT_S_SDE") + SL("dimT_VS_SDE")
    SL("T4C_D") = T_sta_FastDE - dimT_SDE '+ remT_WSstart
    SL("T_STA_slowDE") = SL("T4C_D")
    T_sta_SlowDE = SL("T4C_D")

  'Phase 4 Remontee S et VS
    If ht_step > 0 Then 'steching
       SL("T4C_Slow") = 0
       SL("T4C") = 0
    Else 'remontée concerne uniqt raccsst
       SL("T4C_Slow") = T_sta_SlowDE + SL("remT_F") + SL("remT_S")
       SL("T4C") = SL("T4C_Slow") + SL("remT_VS") ' =T0c theoriqt c'est une vérif
    End If

    T_STA_S = SL("T4C_Slow") ' =  T_sta_SlowDE + SL("remT_F") + SL("remT_S")

    SL("T_max") = T_sta_FastDE + remT_WSstart - dimT_SDE
    T_MAX = SL("T_max")

 SL.Update

End Sub
```

#### Sub AA_REMontees_trace_D_stepl_OK2() ' tracé des exponenteilles

```
e_tt = 8 'mis ds aa_remontees trace tout au debu't

O.DrawWidth = 3
If Fresc.Tag = 6 And REMr2.RecordCount > 0 Then
    REMr2.Index = "PK"
    dbM.Execute "DELETE * FROM REM_r2 ;"
End If

'EVt Fast   de T_sta_Vi à T_STA_F ..... dépend de p_startF
   ‘t_preF = 0   'to_startF = 0.6/0.7ms

'EVt Slow  de T_STA_F à T_STA_S ........dépend de p_startS et et p_sartH         'mais pour choix_TOUT on partira de T_sta_slowDE (T4C_D)
   to2_startS = 0

'EVt Very Slow   de T_STA_S à  T_Max  ....  dépend de (1-p_startF-p_startS) et (1-p_sartH)   'mais pour choix_TOUT on partira de
T_sta_slowDE (T4C_D)
   to2_startVS = 0

'EVt FastDE  de T2C_Vi0 à T3C_A.... double exponentiellep our les détachments afin d'amortir ???

'EVt SlowDE  de T3C-A à T4C_D ....'double exponentiellep our les détachments afin d'amortir ???

'T_max = REMo("T_max") = REMo("T4C") = REMo("T4C_Slow") + REMo("remT4_VS") ' =T0c theoriqt c'est une vérif

If Frpres.Tag = 0 Or Frpres.Tag = 2 Then 'légendes
    O.DrawWidth = 3
    xmi = -xn * 100 / (xx - xn)
    ymi = T_deb * 100 / (yx - yn)
    If Frtyp.Tag < 3 Then 'mode L = croix
        O.CurrentX = xmi - 0.5: O.CurrentY = ymi + 1: O.Line -Step(1, -2), coul_2 ', BF
        O.CurrentX = xmi + 0.5: O.CurrentY = ymi + 1: O.Line -Step(-1, -2), coul_2
    Else 'mode A = carre
        O.CurrentX = xmi - 0.5: O.CurrentY = ymi + 1: O.Line -Step(1, 0), coul_2 ', BF
        O.CurrentX = xmi + 0.5: O.CurrentY = ymi + 1: O.Line -Step(0, -2), coul_2 ', BF
        O.CurrentX = xmi + 0.5: O.CurrentY = ymi - 1: O.Line -Step(-1, 0), coul_2
        O.CurrentX = xmi - 0.5: O.CurrentY = ymi - 1: O.Line -Step(0, 2), coul_2
    End If
    O.ForeColor = coul_2
    z = Int(T_deb + 0.5)
    O.CurrentX = xmi - (4 + Len(z)): O.CurrentY = ymi + 1: O.Print "T1C=" + Format(z)
End If

 O.DrawWidth = e_tt
'                        eVi   eF      eS       eVS      eFDE         eSDE       TOUT
 xmi = Choose(Fresc.Tag + 1, 0, t_preF, t_preS, t_preVS, t_pre_FastDE, t_pre_SlowDE, 0)
 '                           eVi     eF              eS                  eVS     eFDE        eSDE       TOUT
 ymi = Choose(Fresc.Tag + 1, T_deb, T_STA_Vi, T_STA_S - REMo("remT_S"), T_STA_S, T_STA_Vi, T_sta_FastDE, T_deb)
O.CurrentX = (xmi - xn) * 100 / (xx - xn)
 O.CurrentY = (ymi - yn) * 100 / (yx - yn)

Npts = xx * 5 '1200
For i = 0 To Npts
     xav = i / 5
    'on calcule "ymi" ds SUB_AA_REMontees_trace_D_stepl_OK2_calculs_pT  puis on l'introduit dans "yav"
     Select Case Fresc.Tag

       Case 0 'VIiscosité de T_deb=T1C(=pT1_Elas+Visc) à T_sta_Vi (=pT1_Elas)
           AA_REMontees_trace_D_stepl_OK2_calculs_pT IIf(Frtyp.Tag < 3, "Relax", "MIX_Relax_remT_F")
           yav = T_deb + ymi

       Case 1 'eF : Fast   de T_STA_Vi à T_STA_F
            AA_REMontees_trace_D_stepl_OK2_calculs_pT "remT_F"
            yav = T_STA_Vi + ymi

       Case 2 'eS 'evt Slow_startWS Slow de T_STA_F  à T_STA_S
            AA_REMontees_trace_D_stepl_OK2_calculs_pT "remT_S"
            yav = T_sta_SlowDE + SL("remT_F") + ymi ' T_STA_S = SL("T4C_Slow") ' =  T_sta_SlowDE + SL("remT_F") + SL("remT_S")
            If ymi = 0 Then xav = -99
```

```
Case 3 'eVS 'evt Very Slow_startWS Slow de T_STA_S à T_MAX=T4C=T0C
     If t_preVS >= xx Then Exit For
    AA_REMontees_trace_D_stepl_OK2_calculs_pT "remT_VS"
     yav = T_STA_S + ymi
     If ymi = 0 Then xav = -99

Case 4 'eFDE : evt FastDE = descente T3du au detachment>teTa3  de T_sta_vi à T_STA_FastDE = REMo("T3C_A")
      AA_REMontees_trace_D_stepl_OK2_calculs_pT "dimT_FDE"
      yav = T_STA_Vi + ymi 'ymi<0
       If ymi = 0 Then xav = -99

Case 5 'eSDE : evt SlowDE = descente T4 du au detachment>teTa2  de T_STA_FastDE=T3C_A à T_STA_slowDE=T4C_D

     AA_REMontees_trace_D_stepl_OK2_calculs_pT "dimT_Te_SDE"
     yav = T_sta_FastDE - ymi '>0
      AA_REMontees_trace_D_stepl_OK2_calculs_pT "dimT_F_SDE"
     yav = yav - ymi
      AA_REMontees_trace_D_stepl_OK2_calculs_pT "dimT_S_SDE"
     yav = yav - ymi
     AA_REMontees_trace_D_stepl_OK2_calculs_pT "dimT_VS_SDE"
     yav = yav - ymi

 Case 6 'tout'
     AA_REMontees_trace_D_stepl_OK2_TOUT
    REMr2.AddNew
      REMr2("cas") = zfm
      REMr2("step") = t
      REMr2("pt") = i
      REMr2("temps") = xav
      REMr2("pTt") = yav
    REMr2.Update
End Select


If xav <> -99 Then
   'c'est ici qu'il faut faire le calcul de regression
   xav = (xav - xn) * 100 / (xx - xn)
   yav = (yav - yn) * 100 / (yx - yn)
   O.Line -(xav, yav), coul ' QBColor(1)
End If

Next 'i= point

End Sub
```

#### Sub AA_REMontees_trace_D_stepl_OK2_calculs_pT(iX As String)
#### ' tracé des exponenteilles

```
 Select Case iX
'*** relaxation depart Viscosité
   Case "Relax" 'de T_deb=T1C(=pT1_Elas+Visc) à T_sta_Vi (=pT1_Elas)
     'ZONE 1 + ZONE 2 ou on la constante t0_Relax augmente pour tenir  compte phenomene de slack ?
     If ht_step > -L_WS Then  'Abs(z1_min)
        ymi = (T_STA_Vi - T_deb) * (1 - Exp(-xav / to_Relax))
     Else
        ymi = 0
     End If

'*** Remontées
  Case "remT_F" 'eF : Fast   de T_STA_Vi à T_STA_F"    'REMo("remT2_mR")=T_STA_F - T_STA_Vi
        t_preF = IIf(Frtyp.Tag = 1 Or Frtyp.Tag = 3, 0, to_Relax)
        If xav >= t_preF Then
           ymi = SL("remT_F") * (1 - Exp(-(xav - t_preF) / to_startF))
        Else
          ymi = 0
        End If

   Case "remT_S" 'eS : Slow de T_STA_F  à T_STA_S"
        If xav >= t_preS Then
             ymi = SL("remT_S") * (1 - Exp(-(xav - t_preS) / to1_startS))
         Else
           ymi = 0
        End If

  Case "remT_VS" 'eVS 'evt Very Slow_startWS Slow de T_STA_S à T_MAX=T4C=T0C
         If xav >= t_preVS Then
           ymi = SL("remT_VS") * (1 - Exp(-(xav - t_preVS) / to1_startVS))
         Else
            ymi = 0
        End If

'**** Fast DE
  Case "dimT_FDE"  'eFDE : evt FastDE = descente T3du au detachment>teTa3  de T_sta_vi à T_STA_FastDE = SL("T3C_A")
        If ht_step > -L_WS And ht_step < X_Z1 And xav >= t_pre_FastDE Then
             ymi = (T_sta_FastDE - T_STA_Vi) * (1 - Exp(-(xav - t_pre_FastDE) / to_FastDE))
        Else
             ymi = 0
        End If

'**** Slow DE
   Case "dimT_Te_SDE" 'eSDE : evt Tetanos suivi par SlowDE = descente T4 du au detachment>
        If ht_step > -L_WS And ht_step < 0 And xav > t_pre_SlowDE Then  '  Y0       +(Ysta-Y0)
          ymi = SL(iX) * (1 - Exp(-(xav - t_pre_SlowDE) / to_SlowDE))
        Else
           ymi = 0
        End If

   Case "dimT_F_SDE" 'eSDE : evt startF suivi par SlowDE = descente T4 du au detachment>teTa2
       D = t_pre_SlowDE + to_startF
      If ht_step < -aX And xav > D Then  '  Y0       +(Ysta-Y0)
         ymi = SL(iX) * (1 - Exp(-(xav - D) / to_SlowDE))
        Else
          ymi = 0
        End If

 Case "dimT_S_SDE"  'eSDE : evt startS suivi par SlowDE = descente T4 du au detachment>teTa2
          D = t_pre_SlowDE + t_preS
           If ht_step < -aX And xav >= D Then
             sX = to1_startS * Exp(-(xav - D) / to1_startS) / (to1_startS - to_SlowDE)
             sXX = to_SlowDE * Exp(-(xav - D) / to_SlowDE) / (to_SlowDE - to1_startS)
             ymi = SL(iX) * (1 - sX - sXX)
          Else
             ymi = 0
          End If
```

```
   Case "dimT_VS_SDE" 'eSDE : evt startVS suivi par SlowDE = descente T4 du au detachment>teTa2
      D = t_pre_SlowDE + t_preVS
        If ht_step < -aX And xav >= D Then
        sX = to1_startVS * Exp(-(xav - D) / to1_startVS) / (to1_startVS - to_SlowDE)
        sXX = to_SlowDE * Exp(-(xav - D) / to_SlowDE) / (to_SlowDE - to1_startVS)
        ymi = SL(iX) * (1 - sX - sXX)
      Else
         ymi = 0
      End If
   Case Else: Stop

End Select

End Sub
```

#### Sub AA_REMontees_trace_D_stepl_OK2_TOUT()

```
    AA_REMontees_trace_D_stepl_OK2_calculs_pT "Relax"
    yav = T_deb + ymi

    AA_REMontees_trace_D_stepl_OK2_calculs_pT "remT_F"
    yav = yav + ymi

    AA_REMontees_trace_D_stepl_OK2_calculs_pT "dimT_FDE"
    yav = yav + ymi '<0

    AA_REMontees_trace_D_stepl_OK2_calculs_pT "dimT_Te_SDE"
    yav = yav - ymi '>0

    AA_REMontees_trace_D_stepl_OK2_calculs_pT "dimT_F_SDE"
    yav = yav - ymi

    AA_REMontees_trace_D_stepl_OK2_calculs_pT "dimT_S_SDE"
    yav = yav - ymi

    AA_REMontees_trace_D_stepl_OK2_calculs_pT "dimT_VS_SDE"
    yav = yav - ymi

    AA_REMontees_trace_D_stepl_OK2_calculs_pT "remT_S"
    yav = yav + ymi

    AA_REMontees_trace_D_stepl_OK2_calculs_pT "remT_VS"
    yav = yav + ymi

End Sub
```

#### CP5.2 Calculation and curve of pT2 as a function of ΔX

The curves on the computer screen are done by calling the "SUB AAPH_ARTNEG_trace()" routine.

Once the data have been recovered (See values displayed in the Tables of Paper 5 and in the Excel sheets of Supplement DA5, the experimental tension from published articles and the theoretical tension curves from model equations are plotted as a function of the time and the hs shortening (Fig CP5.1).

A regression is performed between experimental and theoretical tensions.

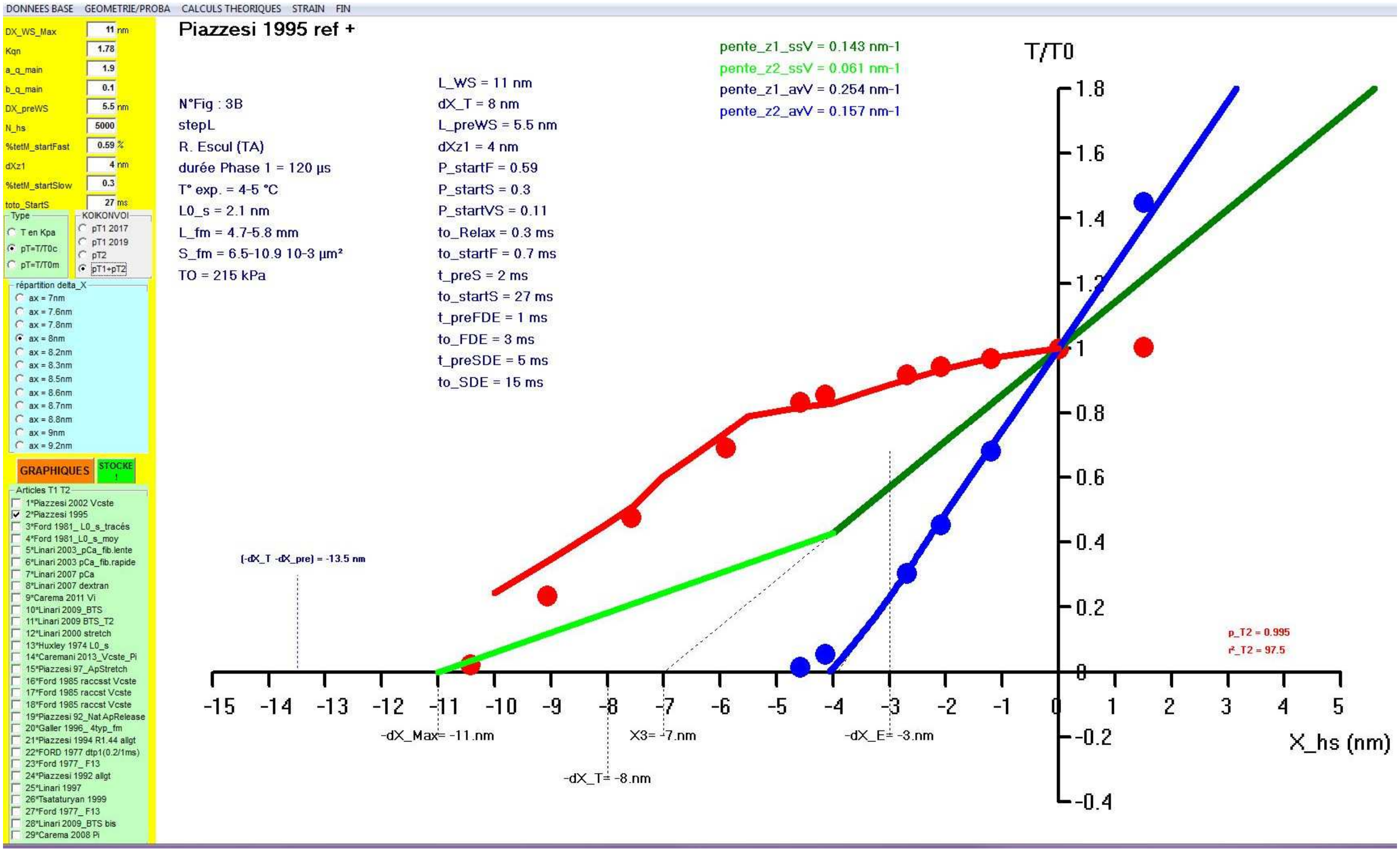


**Fig CP5.2 Screenshot after starting the the "Sub AAPH_ARTNEG_trace()" routine.**

#### Sub AAPH_ARTNEG_trace() *(already seen in Supplement CP4 of Paper 4)*

**….**

```
  T1T2.MoveNext
     Do While T1T2("cas") = z '10000 'z
      If Frpres.Tag <> 2 Then

        'pT2 ***
        If Fresc.Tag > 1 And IsNull(T1T2("X2")) = 0 And IsNull(T1T2("pT2")) = 0 Then
         'If z = 1 Or z = 3 Then coul = QBColor(T1T2("cod_t2"))
         xav = T1T2("X2"): yav = T1T2("pT2") ' / 100
         xav = (xav - xn) * 100 / (xx - xn): yav = (yav - yn) * 100 / (yx - yn)
         If choix_coul = 2 Then
           coul = bcol(T1T2("coul_pt2"))
           O.PSet (xav, yav), coul
         Else
           O.PSet (xav, yav), QBColor(T1T2("coul_pt")) 'special pour T2 QBColor(T1T2("coul_pt2")) por avoir les pts 1995 et 1992
         End If
        End If
       '**********************
       End If
       T1T2.MoveNext: If T1T2.EOF Then Exit Do
     Loop

'PT2 ****** ICI *****
If Fresc.Tag > 1 Then
     F = 1
     If N_fm = 1 Then 'on trace l
       T1T2.Seek "=", zfm, -99
       If IsNull(T1T2("cod_T2")) Then Exit Sub
       If T1T2("cod_T2") = 0 Then Exit Sub
        AAPH_ARTNEG_pT2_OK

     '*** T2 ------------------------------------------ reg T2
      T1T2.Seek "=", zfm, 1: If T1T2.NoMatch Then Stop
      T0_CC = T0c
      Do While T1T2("cas") = zfm
        If IsNull(T1T2("cod_T2")) = 0 Then
          nSM = T1T2("cod_T2"): coul = QBColor(T1T2("coul_pt"))
          Exit Do
        End If
        T1T2.MoveNext
      Loop
      If T1T2("cas") <> zfm Then Stop 'petit verif
      sN = 0: sX = 0: sXX = 0: sY = 0: sYY = 0: sXY = 0
      Do While T1T2("cas") = zfm
        If T1T2("cod_T2") > nSM Then 'on chnde droite
          AAPH_ARTNEG_trace_R2 "T2"
          T0_CC = IIf(Frtyp.Tag = 1, T0(nSM), T0(nSM) / T0(0))
          sN = 0: sX = 0: sXX = 0: sY = 0: sYY = 0: sXY = 0
          nSM = T1T2("cod_T2"): coul = QBColor(T1T2("coul_pt"))
        End If

        If IsNull(T1T2("X2")) = 0 And IsNull(T1T2("pT2")) = 0 And IsNull(T1T2("pT2t")) = 0 Then
          If nSM > 1 Then 'on met à jour
           yap = T1T2("pT2t") * T0(nSM - 1) / T0(0)
           T1T2.Edit: T1T2("pT2t") = yap: T1T2.Update
          End If
          'If T1T2("X2") <= 0 And T1T2("X2") >= -5 Then "espece =1 Rana Esculenta
          If T1T2("X2") <= 0 And T1T2("X2") >= -9 Then  'espece > 1 lesautres Rana T,rat,homme ?
            yav = T1T2("pT2")
            yap = T1T2("pT2t")  'en fait pT2t independant de la tnesion isometrique tetanique
            sN = sN + 1
            sX = sX + yav: sXX = sXX + yav ^ 2
            sY = sY + yap: sYY = sYY + yap ^ 2
            sXY = sXY + yav * yap
          End If
        End If
        T1T2.MoveNext: If T1T2.EOF Then Exit Do
      Loop
  AAPH_ARTNEG_trace_R2 "T2"
```

```
    End If
  End If

End With


End Sub
```

**Sub AAPH_ARTNEG_pT2_OK2(iX As Integer, Icoul As Long)  'Ix =no_courbe  T0ref=1**
**CaLcuLs_Phase2 iX**

```
'on stocke dans T1T2
i = T1T2("cas"): k = T1T2("pt")
T1T2.Seek "=", zfm, 1
sN = 0
Do While T1T2("cas") = zfm
   If T1T2("cod_T2") = iX And T1T2("X2") <= 0 Then
      If T1T2("X2") = 0 Then
        T1T2.Edit: T1T2("pT2t") = 1: T1T2.Update
      Else
         xmi = T1T2("X2")
         z = Int(xmi / 0.5)
         Ph2.Seek "=", zfm, -4: If Ph2.NoMatch Then Stop

         If Ph2("T" + Format(Abs(z))) < 11 Or zfm = 20 Then 'duree phase 2 <11 ms 'on interpole ds table REMP2
            Ph2.Seek "=", zfm, -2: If Ph2.NoMatch Then Stop
            xav = 0.5 * z: yav = Ph2("T" + Format(Abs(z)))
            xap = 0.5 * (z + 1): yap = Ph2("T" + Format(Abs(z + 1)))
            ymi = yav + (xmi - xav) * (yap - yav) / (xap - xav)

            T1T2.Edit: T1T2("pT2t") = ymi: T1T2.Update

         ElseIf xmi >= -8 Then 'cas particilier on garde et on prend valeur du précédent
             T1T2.Edit: T1T2("pT2t") = ymi: T1T2.Update

         Else 'valeur nulle
             T1T2.Edit: T1T2("pT2t") = Null: T1T2.Update
         End If
      End If
   End If
   T1T2.MoveNext
   If T1T2("cod_T2") > iX Then Exit Do
Loop

'on se repositionne
T1T2.Seek "=", i, k

'on trace
Select Case Frtyp.Tag
 Case 1: T0c = T0(iX - 1)
 Case 2: T0c = T0(iX - 1) / T0(0)
 Case 3: T0c = 1
End Select

With O
      .DrawWidth = 8 '5 '8
      .DrawStyle = 0

       'pt de dépat (dx=0;T=T0 ou pT=1)
          .CurrentX = (0 - xn) * 100 / (xx - xn)
          .CurrentY = (T0c - yn) * 100 / (yx - yn)

        'points phase 2
        For k = 1 To tp2_yx * 2 ' nm step de 1 à 15
             Ph2.Seek "=", zfm, -6 'abscisse DX
             xap = Ph2("T" + Format(k))

             Ph2.Seek "=", zfm, -4 'tps_ph2
             xav = Ph2("T" + Format(k))

            If xav < 10 Then
             Ph2.Seek "=", zfm, -2 ' pT2
             yap = Ph2("T" + Format(k))
             Select Case Frtyp.Tag
                Case 1: T0c = T0(iX - 1) * yap
                Case 2: yap = yap * T0(iX - 1) / T0(0) 'T0c = T0(Ix - 1) / T0(0)
                Case 3 ': yap = yap / T0(Ix - 1)
             End Select
```

```
          xap = (xap - xn) * 100 / (xx - xn)
          yap = (yap - yn) * 100 / (yx - yn)
          O.Line -(xap, yap), QBColor(12)

        End If
      Next 'k

     'on trace ligne -dXsatart_dXpre
          .DrawWidth = 1
          .DrawStyle = 2
          xav = -aX - L_preWS: yav = 0: yap = 0.3
          xav = (xav - xn) * 100 / (xx - xn): yav = (yav - yn) * 100 / (yx - yn): yap = (yap - yn) * 100 / (yx - yn)
          O.Line (xav, yav)-(xav, yap)
          .CurrentX = xav - 5
          .CurrentY = yap + 3
          .FontSize = 8
          O.Print "(-dX_T -dX_pre) = " + Format(-aX - L_preWS) + " nm"
          .DrawStyle = 0

End With
End Sub
```

#### Sub CaLcuLs_Phase2(I_courbe As Integer)

```
T1T2met.Seek "=", zfm: If T1T2met.NoMatch Then Stop

 L_WS = T1T2met("L_WS")
 aX = T1T2met("dX_T")
 L_preWS = T1T2met("L_preWS")
 dXz1 = T1T2met("dXz1")
 KqnZ1 = T1T2met("KqnZ1")
 N_hs = T1T2met("N_hs")
to_Relax = T1T2met("to_Relax")
 P_startF = T1T2met("P_startF")
  t_preF = 0 'T1T2met("t_preS")
  to_startF = T1T2met("to_startF")
  P_startS = T1T2met("P_startS")
  t_preS = 0 'T1T2met("t_preS")
  to1_startS = T1T2met("to1_startS")
  t_preVS = T1T2met("to_preVS")
  to1_startVS = T1T2met("to1_startVS")
  t_pre_FastDE = T1T2met("to_preFDE")
  to_FastDE = T1T2met("to_FDE")
  t_pre_SlowDE = T1T2met("to_preSDE")
  to_SlowDE = T1T2met("to_SDE")
P_startH = P_startF + P_startS: If P_startH > 1 Then Stop

'PREPA_calculs
 Set Ph2 = dbM.OpenRecordset("REM_phase2", dbOpenTable): Ph2.Index = "PK"
 If Ph2.RecordCount > 0 Then dbM.Execute "DELETE * FROM REM_phase2"

 Set SL = dbM.OpenRecordset("REMgoodluck_T", dbOpenTable): SL.Index = "PK"
 If SL.RecordCount > 0 Then dbM.Execute "DELETE * FROM REMgoodluck_T"

tp2_xx = 15 'ms
tp2_xt = 0.2 ' ms
Npts = tp2_xx / tp2_xt '5ms / 0.2ms = 25 pts
tp2_yx = 14 'nm ht_step

For t = -6 To Npts 'ms  -1 T2 '-2 tps_T2 -3 ht_step
    xav = t * IIf(t < 1, 1, tp2_xt)
    Ph2.AddNew
      Ph2("fm") = zfm
      Ph2("tps") = xav
   Ph2.Update
Next

Ph2.Seek "=", zfm, -6: If Ph2.NoMatch Then Stop
Ph2.Edit
  Ph2("vu1") = "ht_step"
  For k = 1 To tp2_yx * 2 'step de 0.5 à 15
      ht_step = -k / 2
      Ph2("T" + Format(k)) = Format(ht_step)
  Next 't
Ph2.Update

Ph2.Seek "=", zfm, -4: If Ph2.NoMatch Then Stop
Ph2.Edit: Ph2("vu1") = "tps_ph2": Ph2.Update
Ph2.Seek "=", zfm, -3: If Ph2.NoMatch Then Stop
Ph2.Edit: Ph2("vu1") = "T2": Ph2.Update
Ph2.Seek "=", zfm, -2: If Ph2.NoMatch Then Stop
Ph2.Edit: Ph2("vu1") = "pT2": Ph2.Update

'CALCULS
For k = 1 To tp2_yx * 2 ' nm step de 1 à 15
     ht_step = -k / 2
      SL.AddNew
         SL("cas") = zfm
         SL("courbe") = k
         SL("stepL") = ht_step
         'SL("stepL_Mdeb") = GL("stepL_Mdeb")
         SL("T0") = T1T2met("T0c")
      SL.Update
```

```
     SL.Seek "=", zfm, k: If SL.NoMatch Then Stop
     AA_REMontees_trace_D_stepL_OK1_prepa

     'boucle temps
     For t = 0 To Npts 'ms 'à t=0 correspnnd T0 ou T1 mais on s'en fout
            xav = t * tp2_xt
            AA_REMontees_trace_D_stepl_OK2_TOUT
                      'etape 2 on enregistre
           Ph2.Seek "=", zfm, xav: If Ph2.NoMatch Then Stop
           Ph2.Edit
               Ph2("T" + Format(k)) = yav
               If t > 0 Then Ph2("diff_T" + Format(k)) = yav - yap
          Ph2.Update
          yap = yav
   Next 't
Next 'k

'points phase 2
For k = 1 To tp2_yx * 2 ' nm step de 1 à 15

    If zfm = 20 Then 'Galler n'importe quoi
        Ph2.Seek "=", zfm, tp2_xt * IIf(k = 1, 3, 2)
    Else
        Ph2.Seek "=", zfm, tp2_xt
    End If
    If Ph2.NoMatch Then Stop 'xt=0.2

    ymi = Ph2("diff_T" + Format(k))
    Ph2.MoveNext
    Do Until Ph2.EOF
       If Ph2("diff_T" + Format(k)) < 2 And Ph2("diff_T" + Format(k)) < ymi And Ph2("diff_T" + Format(k)) > 0 Then
          xav = Ph2("tps")
          yav = Ph2("T" + Format(k))
          ymi = Ph2("diff_T" + Format(k))
       End If

       If Ph2("diff_T" + Format(k)) < 0.07 Or (Ph2("diff_T" + Format(k)) > ymi And Ph2("diff_T" + Format(k)) < 1.6) Then
          Exit Do
       End If
      Ph2.MoveNext
    Loop

    'on stocke
          Ph2.Seek "=", zfm, -4 'tps_ph2
          Ph2.Edit: Ph2("T" + Format(k)) = xav: Ph2.Update
          Ph2.Seek "=", zfm, -3 'T2
          Ph2.Edit: Ph2("T" + Format(k)) = yav: Ph2.Update
          Ph2.Seek "=", zfm, -2 'pT2
          Ph2.Edit: Ph2("T" + Format(k)) = yav / T_MAX: Ph2.Update
         'on se repalce
         Ph2.Seek "=", zfm, xav 'tps_ph2
         If Ph2.NoMatch Then Stop
         Ph2.Edit: Ph2("vu" + Format(k)) = IIf(k = 1, "1", 1): Ph2.Update

Next 'k
End Sub
```

#### Sub AAPH_ARTNEG_trace_R2(iX As String)

```
    p_regL = sXY / sXX
    r_regL = 1 - (sYY - 2 * p_regL * sXY + sXX * (p_regL ^ 2)) / (sYY - sY * sY / sN)

    O.CurrentX = 90: O.CurrentY = nSM * 7 + 17.5: O.Print "p_" + iX + " = " + Format(p_regL, "0.###")
    O.CurrentX = 90: O.CurrentY = nSM * 7 + 15: O.Print "r²_" + iX + " = " + Format(r_regL * 100, "###.##")
End Sub
```
